# Generative Landscapes and Dynamics to Design Multidomain Artificial Transmembrane Transporters

**DOI:** 10.1101/2025.03.28.645293

**Authors:** Fernando Montalvillo Ortega, Fariha Hossain, Vladimir V. Volobouev, Gabriele Meloni, Hedieh Torabifard, Faruck Morcos

## Abstract

Protein design is challenging as it requires simultaneous consideration of interconnected factors, such as fold, dynamics, and function. These evolutionary constraints are encoded in protein sequences and can be learned through the latent generative landscape (LGL) framework to predict functional sequences by leveraging evolutionary patterns, enabling exploration of uncharted sequence space. By simulating designed proteins through molecular dynamics (MD), we gain deeper insights into the interdependencies governing structure and dynamics. We present a synergized workflow combining LGL with MD and biochemical characterization, allowing us to explore the sequence space effectively. This approach has been applied to design and characterize two artificial multidomain ATP-driven transmembrane copper transporters, with native-like functionality. This integrative approach proved effective in unraveling the intricate relationships between sequence, structure, and function.

---

Protein design is becoming a cornerstone of modern biotechnology, with possible applications spanning from therapeutic drug design to industrial biocatalysis. Despite its potential, the immense combinatorial space of possible sequences and the challenge of preserving crucial interactions for functional and structural stability make the design process time-consuming and computationally challenging. However, recent advances in machine learning (ML) along with increasing sequence data availability have revolutionized our ability to model biological systems from evolutionary clues. Current applications range from protein structural prediction to the design of novel proteins with enhanced functionality (such as modifying antibody binding affinity to their respective targets or improving catalytic capacity of enzymes) and stability (*1–10*).

Researchers have been exploring diverse ML strategies in protein design, ranging from sequence-based and sequence-labeling models to structure-based and hybrid approaches (*11, 12*). Among these techniques, latent generative models, such as Variational Autoencoders (VAEs), have been only recently explored in the context of biological systems (*3–5, 13*). We introduce a framework that learns from extant protein sequence data and harnesses the reconstructive ability of VAEs in conjunction with Direct Coupling Analysis (DCA) to produce maps of generated sequences known as latent generative landscapes (LGLs) (*14*). These landscapes model protein sequence-function relationships, capturing sequence diversity within a family while assessing functional integrity, offering a powerful tool for protein design.

When designing proteins, it is important to preserve crucial motifs essential for protein function, as well as highly coupled interacting residues that form interdependent networks critical for the protein’s behavior, even beyond conserved sites. A key strength of the LGL framework lies in its ability to learn these key interactions from only protein-family sequences, scoring generated sequence variants more favorably (greater negative value) when such interactions are preserved (*14*). However, relying solely on sequence-based approaches can limit our understanding of intricate interdependencies within proteins, particularly in complex membrane-embedded nanomachines in which catalysis and multidomain coupling are central to protein function. Coupling this method with molecular dynamics (MD) simulations allows for a deeper exploration of how interacting residues and novel mutations influence the protein’s conformational dynamics, domain crosstalk, and functional behavior. This integrated computational approach can bridge the gap between prediction and experimental success. Rather than conducting experiments on numerous designed sequences to identify functional ones, it enables us to achieve remarkable success with a carefully selected subset. This, in turn, allows for dedicating more resources to in-depth validation and characterization of the most promising candidates, an aspect often lacking in currently available protein design studies.

The P-type ATPase superfamily of transmembrane transporters is crucial for cellular homeostasis, coupling ATP hydrolysis to substrate transport across membranes against their electrochemical gradient (*15, 16*). These large, multidomain transmembrane proteins rely on intricate crosstalk between multiple cytosolic and transmembrane domains. Their mechanism couples ATP hydrolysis with auto(de)phosphorylation to trigger dramatic and sequential conformational rearrangements in the cytosolic domains. This results in transmembrane helices movements, which in turn facilitate substrate translocation across the membrane lipid bilayer (*17, 18*). Within this superfamily, the P_1B_-type ATPase subfamily plays an essential role in maintaining transition metal homeostasis, where the P_1B-1_-type ATPases stand out as the ubiquitous Cu(I) exporters in all organisms (*19*). Thus, the complexity, functional diversity, and relevance of this system make it an ideal candidate to test the robustness of the synergized protein design strategy.

Here, we apply an interdisciplinary approach to efficiently design artificial P_1B-1_-type ATPase transporters that maintain their quintessential coupled multidomain complexity and multifunctional properties. First, we employ the LGL framework to produce novel variants of the transporter family, decoding few from the vast sequence space based on favorable Hamiltonian metric and the presence of key subgroup-specific motifs essential for function. Next, we leverage MD to assess the structural stability and dynamics adherence to the expected catalytic scheme. For these generated variants, we demonstrate coupled domain movements crucial for catalytic function, comparable to those identified in the wild-type (WT) protein(s). Finally, we experimentally characterize the preservation of multifunctional properties encompassing membrane embedding, proper fold, copper translocation, and *in vivo* cellular protection from copper toxicity. Our design of complex transmembrane transporters that integrate seamlessly into a lipid bilayer and maintain full functional fidelity advances previous attempts in this field (*20–23*). The rate of success on the selected designs is notable, something rare for sequences with hundreds of mutations, opening new possibilities for creating tailored proteins with specific functional characteristics through an efficient, sequence-driven approach synergized with molecular dynamics.

## Latent generative landscape produces non-extant transporter sequences

The LGL framework enables visual and quantitative exploration of the sequence energy landscape, enhancing mutational analysis, the study of phylogenetic relationships, and identification of different functional clusters (*14*). However, the generative ability to produce complex new protein sequences with desired features has been less explored. The incorporation of the Hamiltonian metric, determined from key interactions learned within the protein subfamily using DCA (see *Methods*), hints at relative functional fitness. This definition of fitness incorporates various aspects of this type of multifunctional transmembrane proteins, such as selective transport of specific substrates, proper energy transduction, coupled conformational changes necessary for activity modulation, and intercommunication between domains. Thus, we hypothesized that the LGL can produce new variants of sequences that preserve these multiple layers of complexity such as the ones present in functional P_1B-1_-type ATPase transporters. These nanomachines couple ATP hydrolysis and auto(de)phosphorylation in the soluble domains, with spatially distant transmembrane substrate translocation via a coupled actuator unit.

The LGL training input consists of a multiple sequence alignment (MSA) of the P_1B_-type ATPase family, comprising a refined set of approximately 13,500 sequences of 574–661 residue length (fig. S1A). This MSA was curated to minimize noise and sampling bias (see *Methods*). Fig. 1A illustrates the methodology used to construct an LGL map of a total of 250,000 pixels, each representing a generated sequence with assigned Hamiltonian values. These generated sequences (pixels) can be decoded and analyzed in synergy with MD simulations, serving as a powerful integrated tool for protein design that effectively narrows the search space, enabling comprehensive characterization of the artificial transporters. When the P_1B_-type ATPase training data is plotted on the LGL in Fig. 1B, a clustering pattern emerges, aligning with the established classification of the subgroups based on reported transmembrane motifs that correlate with transition metal cargo selectivities (*18*). Though the LGL allows us to potentially sample thousands of generated sequences within the subfamily, we narrowed down the selection to the P_1B-1_ subgroup. This highly characterized subgroup, with available crystal structures, enables a robust and meaningful comparison and plays essential physiological roles across all organisms, including humans (*16, 18, 24*). Since P_1B-1_ sequences from the training dataset appear in the lower half of the map as purple symbols in Fig. 1B, we hypothesize that the sequences decoded from this region are more likely to have P_1B-1_-like functionality. With this in mind, we focused on the area that contained the well-characterized Cu(I)-pump from *Legionella pneumophila* (*Lp*CopA) sequence for which structural information is available (PDB: 3RFU), thus serving as a reference. Fig. 1C depicts a 3D representation of the region, where the red circle represents the location of WT *Lp*CopA sequence, and the nearby white circles, two decoded generated sequences GS1 and GS2 (selected out of eight characterized by MD, *vide infra*). Additionally, we ensured that the decoded sequences did not directly overlap with existing native sequences to increase the diversity of novel mutations (fig. S1B).

**Figure 1:**
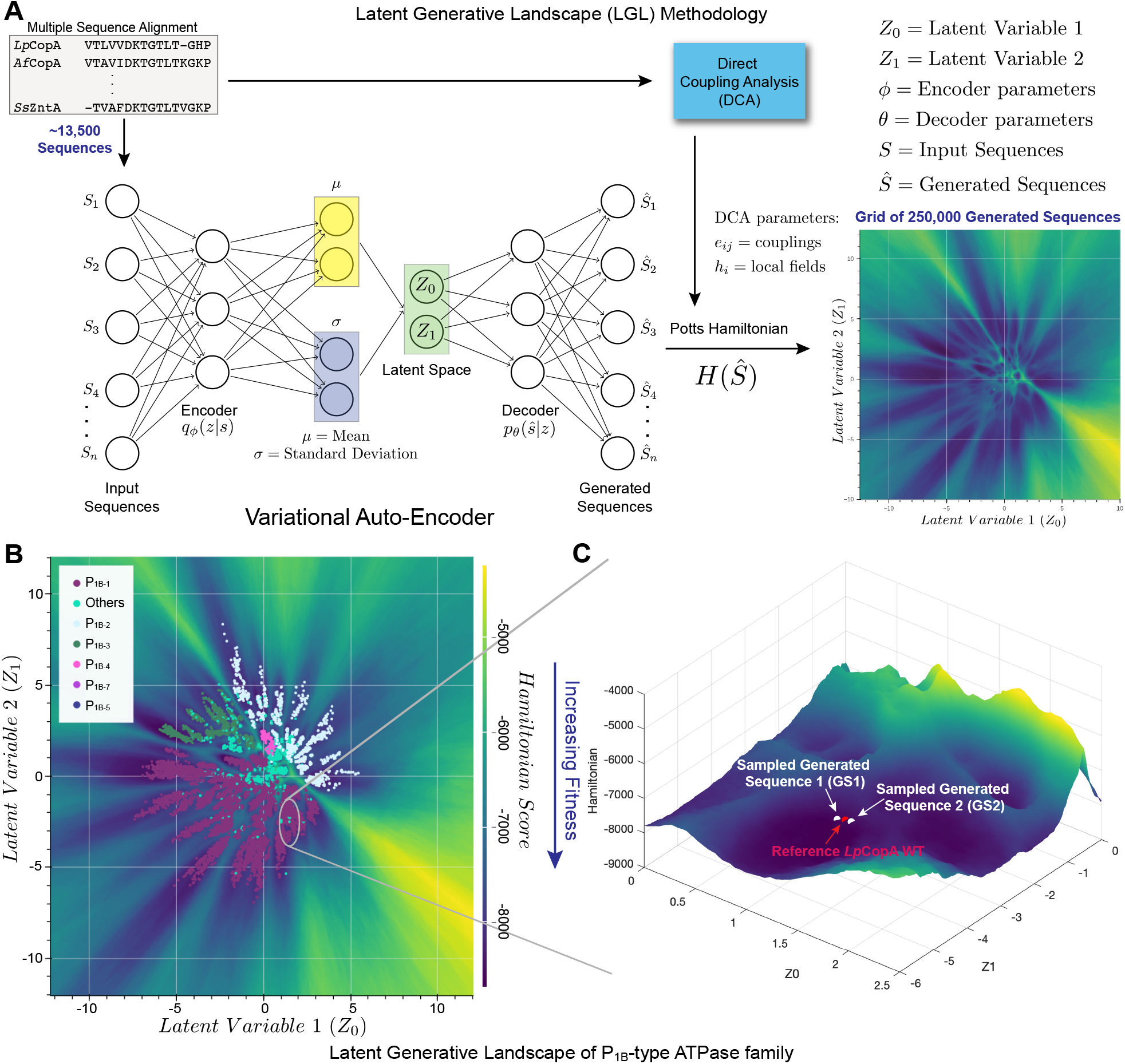
Overview of the Latent Generative Landscape (LGL) methodology. (**A**) The algorithm combines the generative power of the Variational Auto-Encoder (VAE) and the Potts Hamiltonian to approximate fitness of the generated sequences, resulting in an LGL map. (**B**) The LGL plot was generated using the assembled MSA for the P_1B_-type ATPase family. Labeled sequences part of the training dataset (13,553 sequences) are plotted on the LGL to identify the corresponding subgroup locations. (**C**) A 3D landscape view of the decoded region with reference *Lp*CopA labeled with a red circle while the two selected decoded sequences (GS1 and GS2) labeled in white.

## Structural stability and mutation distribution of generated sequences

The domain topology diagram in Fig. 2A illustrates that P_1B-1_-type ATPases consist of a trans-membrane (TM) domain featuring eight TM helices (MA-M6) responsible for metal substrate binding and forming the translocation pathway, three cytosolic domains (N-, nucleotide binding; P-, phosphorylation; A-, actuator), and metal-binding domain(s) (MBDs), located at either the N or C-terminal, serving a regulatory function (*25*). The deletion of MBDs has been shown to reduce the transport rate without affecting the transport mechanism or selectivity (*26, 27*). Therefore, it was excluded from our LGL training dataset and subsequent analysis.

**Figure 2:**
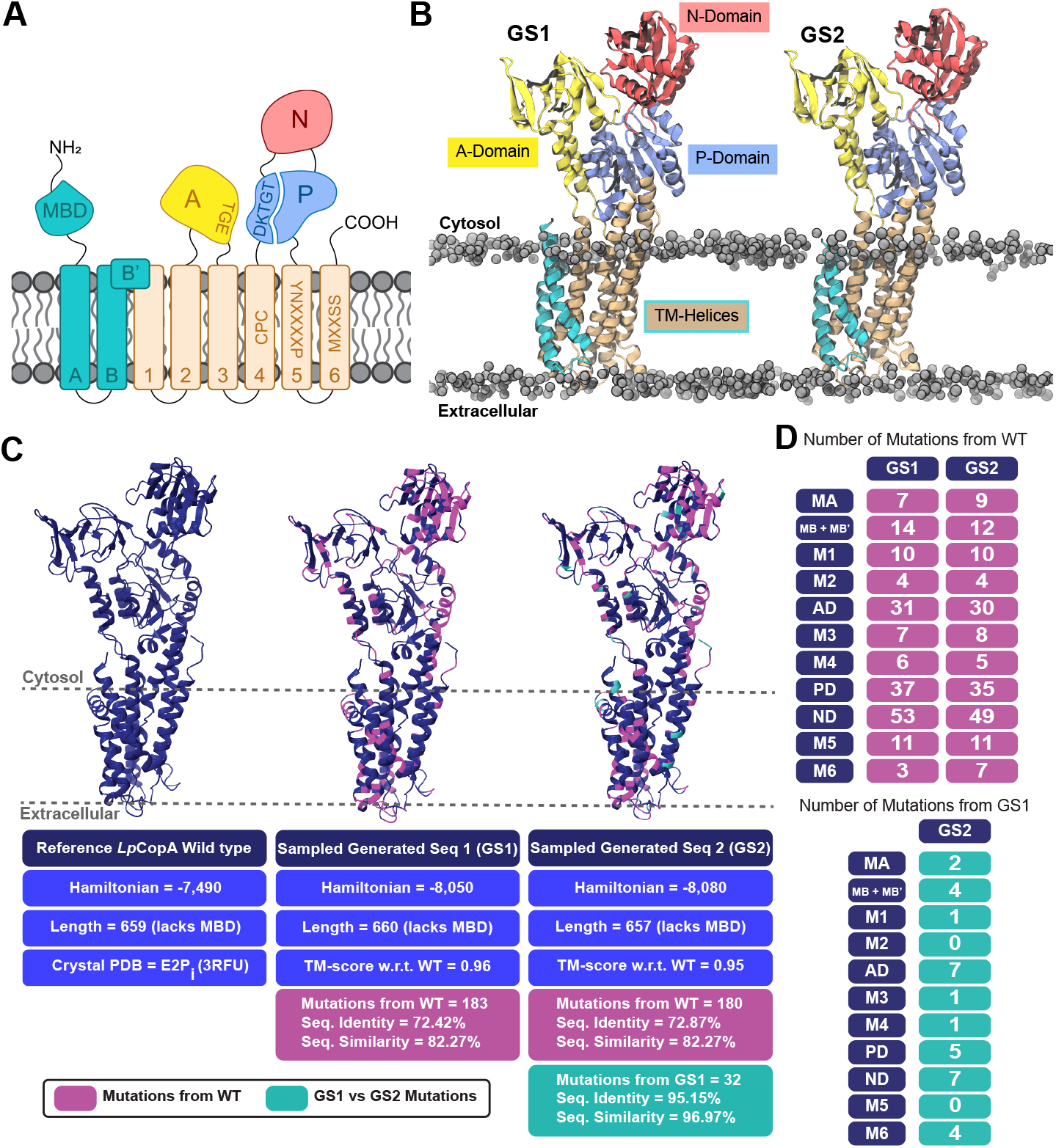
Structural and sequence composition details of the P_1B-1_-type ATPase subgroup and the two selected decoded generated sequences. (**A**) A 2D domain topology of the P_1B_-type ATPase family with highlighted conserved motifs. DKTGT and TGE motifs on the P-domain and A-domain, respectively, are key in the auto(de)phosphorylation catalytic cycle characteristic of P-type ATPase transporters. Specific substrate selectivity motifs associated with the P_1B-1_-type ATPase subgroup are labeled on M4, M5, and M6. (**B**) A 3D domain topology and membrane insertion of the generated proteins in lipid bilayers, pre-screened via the PPM 2.0 server for membrane compatibility prior to MD simulation. (**C**) Comparison of structural, sequential, and fitness characteristics of GS1 and GS2 relative to WT *Lp*CopA. TM-score values indicate that the AF2-predicted structures closely match the crystal structure. Magenta highlights mutations relative to WT, while teal shows mutations relative to each other between the generated sequences. The Hamiltonian values infer that the generated sequences would maintain native-like functional fitness when compared to *Lp*CopA. (**D**) Domain-specific mutation count comparison for GS1 and GS2 versus *Lp*CopA in the top table, with the bottom table showing mutation counts between GS1 and GS2.

To better understand the distribution of the novel mutations within the described topological framework, Fig. 2C highlights the mutation locations in the generated sequences, with corresponding labeled sequence alignments provided in fig. S2 and fig. S3. To explore the effect of these mutations on the 3D structure and stability, the sequences were first analyzed using TOPCONS to predict membrane topology, number and location of transmembrane helices, and delineate intracellular and extracellular domains (*28*). The predicted domain topology included eight TM helices and two large intracellular soluble regions that closely resemble that of the WT (fig. S4A). Consequently, AlphaFold2 (AF2) and Positioning of Proteins in Membranes 2.0 (PPM2.0) were employed to obtain structural and membrane insertion predictions for GS1 and GS2 presented in Fig. 2B (*1, 29*). The structures obtained showcase the characteristic eight TM helices and three catalytic soluble domains, as well as correct lipid bilayer insertion. The sequences had template modeling scores near 1 as reported in Fig. 2C (*1, 26*). This metric is commonly used in structural biology to quantify fold resemblance, with values ranging from 0 to 1, where 1 indicates a perfect match and an identical fold to the WT protein (*30*).

Relative to *Lp*CopA, both GS1 and GS2 contained approximately 180 novel mutations, despite sharing 70% identity to the WT. These mutations were distributed throughout the protein where approximately 24% of the A-domain, 26% of the P-domain, 42% of the N-domain, and 23% of the TM region were mutated (Fig. 2D). This result implies that the VAE provides each domain with flexibility in accommodating mutations without significantly disrupting the overall structure. A closer inspection revealed that both sequences contained a single insertion at V365 in the N-domain compared to the WT. This was notable, as removing this residue caused distortion of the first N-domain *β*-sheet (P359 to A367 in GS1 and P358 to A366 in GS2) in the AF2 model (fig. S4B). Further MD simulations showed that the secondary structure was not recovered after 400 ns (fig. S4C). Interestingly, this insertion aligns appropriately with other P_1B-1_ sequences (see position 397 in fig. S3), highlighting the predictive power of our generative model. Additionally, the generated sequences have 32 mutations among themselves with mutations scattered throughout the protein (Fig. 2, C and D). Despite these variations, the Hamiltonian values reported for each sequence in Fig. 2C are comparable, suggesting that these sequences maintain similar functional properties.

## Molecular dynamics identifies crucial dynamics of generated transporters

With high confidence in the structural integrity of the generated sequences, we performed MD simulations to investigate crucial dynamical properties. Structural characterizations of CopA in both eukaryotic and prokaryotic systems reveal a high degree of similarity (PDB: 3RFU, 8Q75, 7SI3, 4BBJ, and 8Q76 represent E2P/E2P_*i*_; PDB: 7R0I, 8Q74, and 7XUM represent E1-copper bound; PDB: 7R0G, 8Q73, and 7XUN represent E1-apo), indicating conservation of the Post-Albers mechanism cycle across the kingdoms (Fig. 3A) (*19, 24, 26, 31–33*). This transport mechanism involves the intricate and coupled interplay of soluble domain movements driven by auto(de)phosphorylation events. These movements induce rearrangements within the TM domain, allowing the protein to alternate between two major states: the E1 inward-facing state, which has high affinity towards the substrate, and the E2 outward-facing state with lower affinity (*34*). Particularly, the E2P_*i*_ state is the starting point of a significant conformational change of the A-domain as the system evolves towards the E1 state (*26, 31*). This A-domain movement triggers a substantial reorganization of the TM domain, visualized as two blocks of four TM helices each, (MA, MB, M1, M2) and (M3, M4, M5, M6), which move relative to each other, opening and closing the translocation pathways and making the binding site accessible to opposite sides of the lipid bilayer (*31*).

**Figure 3:**
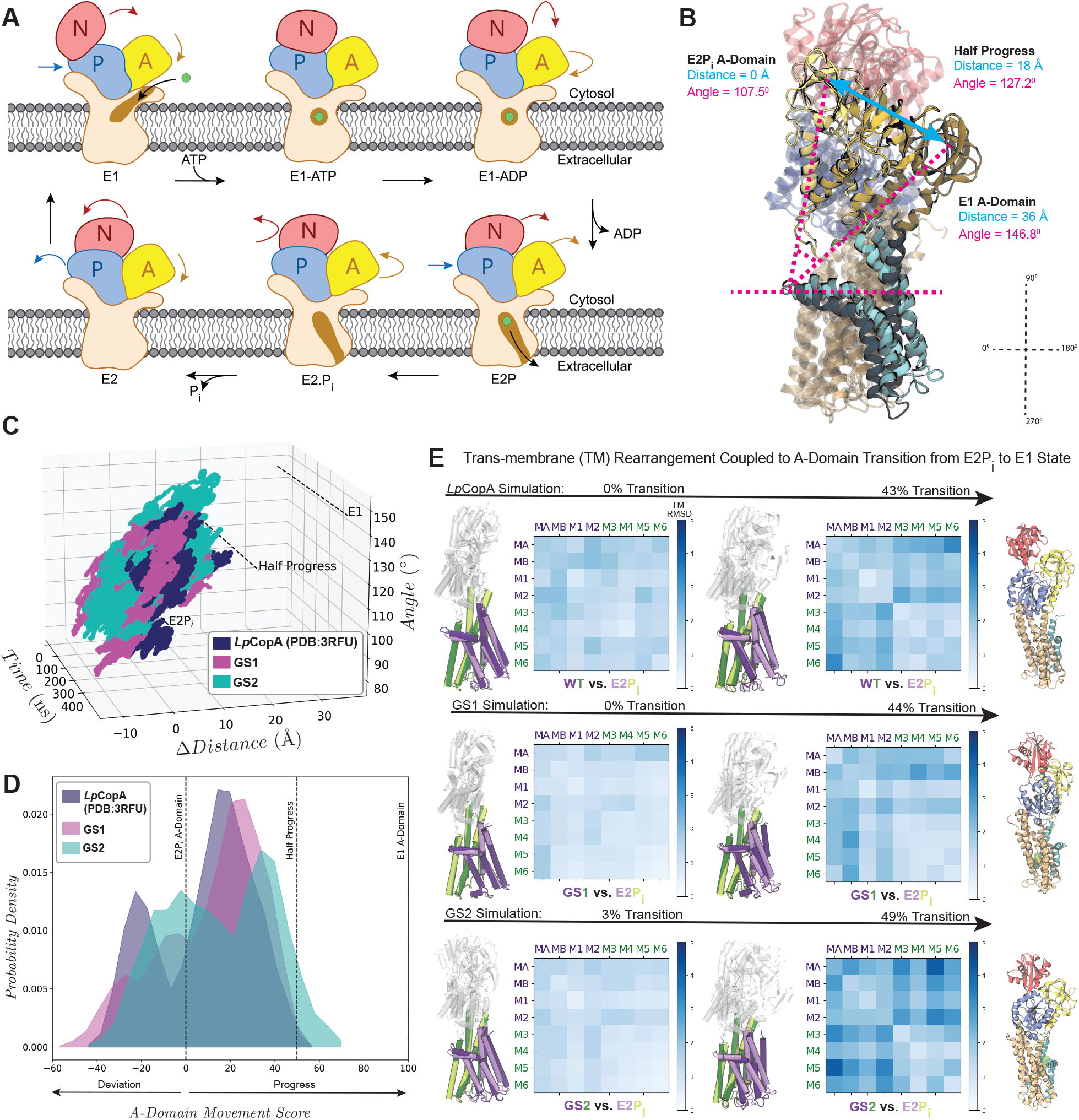
Molecular dynamics to capture key mechanisms and domain coupling required for metal substrate transport. (**A**) The P_1B_-type ATPase family follows the Post-Albers cycle. The pump converts between two states, E1 and E2, in an alternating access mechanism. Metal(s) bind to TM site(s) (E1 state), are occluded within the membrane upon ATP hydrolysis/phosphorylation (E1P), and released on the opposite side in the E2P state, followed by a dephosphorylation transition state (E2P_*β*_) to then regenerate E1. During the E2P_*β*_ to the E1 transition, the A-domain undergoes a significant transition that is coupled to a rearrangement of the transmembrane transport pathway. (**B**) Structural alignment of the different stages of the A-domain transition along with labeled vectors used to calculate the tilt angle and ωdistance. (**C**) A-domain dynamics were measured over 400 ns simulations by tracking changes in tilt angle and ωdistance for *Lp*CopA, GS1, and GS2. (**D**) The *A-domain movement scores* for GS1 and GS2 show transitions similar to the WT. (**E**) Transmembrane helices rearrangement panels consist of structural alignments and TM helix distance difference matrices in relation to labeled *A-domain movement scores*. The purple block (MA-M2) and green block (M3-M6) represent the regions of the protein, with the E2P_*β*_ state shown in lighter tones and later A-domain stages in darker tones. The top row illustrates the rearrangement in the WT *Lp*CopA simulation followed by GS1 and GS2, respectively. The last column displays CAVER tunnels for each.

Our initial decoded list comprised of eight generated sequences (GS1-GS8) with varying Hamiltonian values and locations on the LGL map (Fig. 2C and fig. S5, A and D). Numerical Hamiltonian value alone, however, does not fully convey the extent to which interactions are retained or altered, nor their impact on the coupled dynamics. We aimed to explore various LGL regions around the *Lp*CopA area to gain a deeper understanding of the relationship between the calculated Hamiltonian and its impact on the system’s dynamics. Integrating MD simulations allowed us to unravel intricate details regarding their A-domain and TM helix bundle rearrangement properties, enabling the evaluation of their consistency with the Post-Albers cycle and available crystal structures.

To assess the viability of the generated sequences as functional candidates, MD studies were conducted for *Lp*CopA and GS1-GS8. Each system underwent several 400 ns trials within an isobaric-isothermal (NPT) ensemble, with the protein embedded in a lipid bilayer and surrounded by a salt-containing aqueous solution. To exemplify the evaluation process, we will focus our discussion on the results of *Lp*CopA (PDB: 3RFU), GS1, and GS2, which were selected due to their native-like dynamics and favorable Hamiltonian values. Fig. 3B illustrates the A-domain progression at different stages of the transition from E2P_*i*_ (PDB: 3RFU) to E1 (PDB: 7R0I) (outward-to inward-open) as well as the two metrics used for its movement quantification: the tilt angle (pink lines) and the ωdistance (blue lines). The distribution of these two values over the simulation time for the three systems is shown in Fig. 3C, highlighting significant dynamics in the A-domain (see Movie S1). The angle and distance values were combined into a one-dimensional metric, termed the *A-domain movement score*, using equations S4-S7. Fig. 3D shows that positive values, defined as “progress”, correspond to A-domain motions toward the E1 state, while negative values, referred to as “deviation”, indicate motions in the opposite direction. Although WT *Lp*CopA did not fully transition to the E1 state due to the limitations associated with capturing large conformational changes using all-atom simulations, the overlapping bimodal distribution among the three proteins unravels similar A-domain dynamics to those of the WT, indicating comparable coupled domain movements characteristic of the Post-Albers catalytic cycle. The results for each system trial are detailed in fig. S6, A and C.

Fig. 3E summarizes the qualitative and quantitative analyses used to assess the TM inter-block rearrangement. The superposition of structures, with purple representing the first TM block and green the second block, was combined with the helix Distance Difference Matrix (DDM) to observe and quantify the rearrangement. Results from the *Lp*CopA simulations (Fig. 3E *top* panel), show a pattern clearly mimicking the proper inter-block distance changes observed in experimentally determined crystal structures in fig. S6B (*19, 26, 31*). In the DDM for the 100% A-domain progress in fig. S6B, a large inter-block rearrangement is observed at the bottom-left and top-right corners, while the intra-block rearrangement showed only small RMSD values. We demonstrated that, as the A-domain of both generated sequences transitions from the E2P_*i*_ to the E1 state, the TM block movement begins to resemble that of a coupled native transporter, with GS2 showing this more clearly (Fig. 3E *bottom* panel, Movie S2). GS2 also starts to exhibit an E1-like transmembrane tunnel, as indicated by CAVER 3.0 analysis (Fig. 3E *right-most bottom* panel) (*35*). These results suggest that the modeled systems retain the essential mechanistic features required for their biological activity, revealing dynamic properties that complement the LGL approach.

The A-domain and corresponding helix bundle movement rearrangement analyses were also carried out for the GS3-GS8 sequences (fig. S5B-D; tables S1-S2; see *Supplemental* for detailed analysis). Synchronizing LGL predictions with MD simulations allowed us to identify which aspects of these dynamics in the decoded eight sequences were impacted and to what extent, a valuable insight in addition to the Hamiltonian values. Notably, GS1 and GS2 demonstrated superior performance in preserving these desired dynamic properties and thus were subjected to experimental characterization.

## Experimental characterization of preserved WT functionality

The two selected generated sequences were subjected to experimental characterization to validate membrane insertion, correct folding, *in vitro* catalytic ATP hydrolysis activity, and *in vivo* cellular protection from copper toxicity underlying transport capability. TM domain folding was assessed by analyzing membrane localization upon recombinant expression in *E. coli* membrane fraction (MF), while folding of the soluble domains was validated through *in vitro* ATP hydrolysis activity, ensuring the protein retained its ATP hydrolysis and P-domain’s autophosphorylation turnover. Copper transport activity was evaluated *in vivo* by testing recombinant GS1 and GS2 expression via intracellular copper quantification and protection from copper toxicity.

Initial results, presented in the Western blot in Fig. 4A, confirmed successful recombinant expression despite each generated protein featuring approximately one-third of its sequence mutated. Both protein constructs were primarily localized to the MF, indicating proper insertion in the lipid bilayer and TM domain folding, further supported by the significantly monodisperse Size Exclusion Chromatography (SEC) profiles upon extraction and purification in detergent micelles (fig. S7A). No significant misfolded proteins were indeed observed in the soluble fraction (SF) or inclusion bodies.

**Figure 4:**
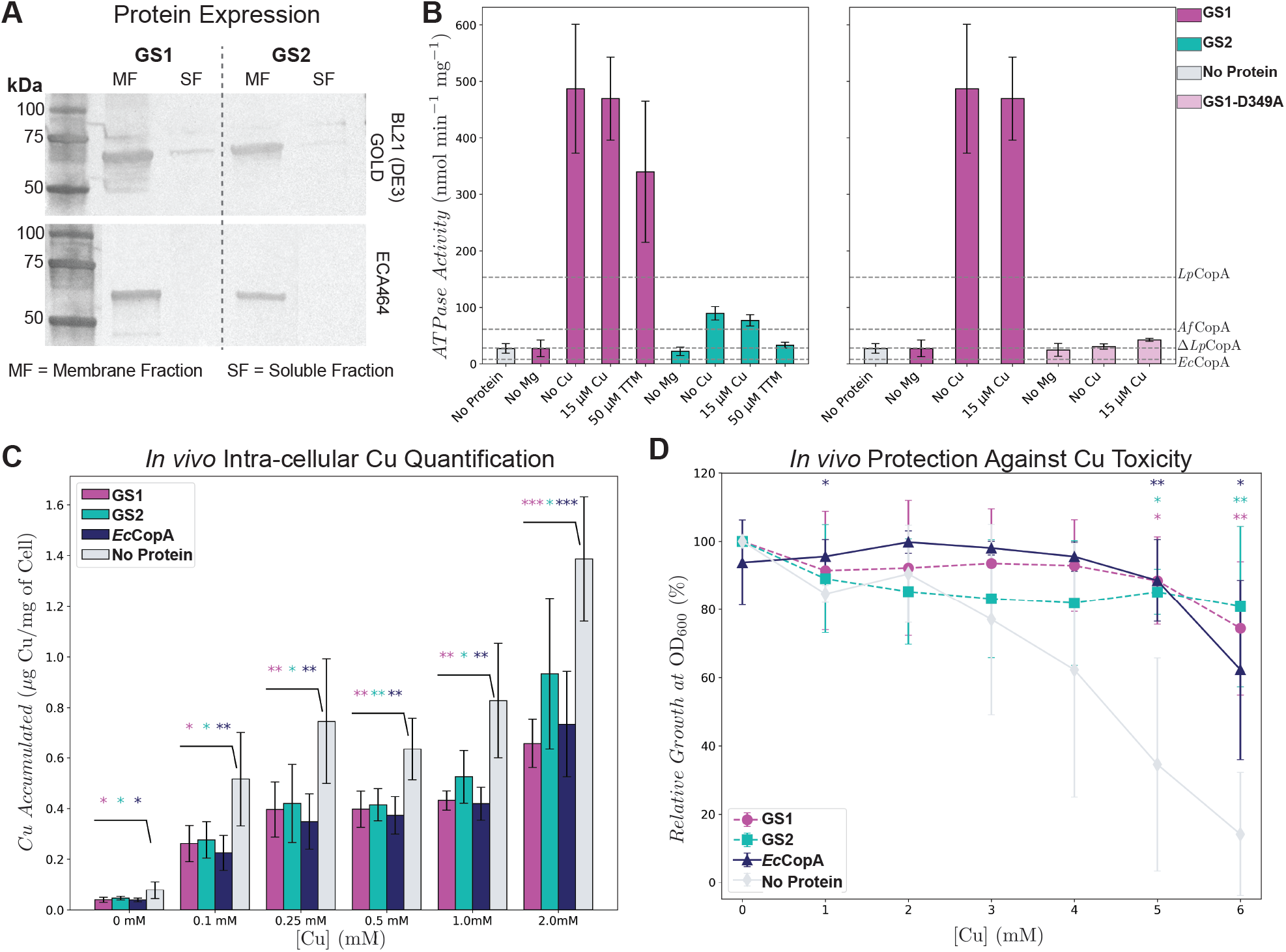
Experimental characterization of expression, membrane embedding, hydrolysis activity, and copper export for GS1 and GS2. (**A**) Western blot analysis reveals successful protein expression and insertion of both GS1 and GS2 in the membrane fraction. (**B**) *In vitro* ATPase activity rate for GS1 and GS2 in comparison to several characterized P_1B-1_ transporters (dashed line). WT ω*Lp*CopA lacks the MBD, providing an ideal comparison with GS1 and GS2. GS2 exhibited rates that are in range with the reported values for characterized prokaryotic P-type ATPases, while GS1 showed a significantly higher catalytic rate (*17, 26, 36*). The GS1-D349A mutation, which abolished ATP hydrolysis activity, confirmed that the defining auto(de)phosphorylation of the aspartate in the conserved DKTGT motif in the P-domain was preserved. (**C**) *In vivo* e#ux assay demonstrates that both GS1 and GS2 can effectively export copper from cells as a function of copper concentrations, similar to WT *Ec*CopA, while the absence of a transporter leads to metal accumulation within the cells. (**D**) *In vivo* copper-susceptibility growth assay confirms that both GS1 and GS2 can protect cells against copper toxicity and promote cell viability as a function of increasing copper concentration, similar to overexpression of WT *Ec*CopA. Statistical significance was calculated by unpaired Student’s *t*-test (*** for p *<*0.001, ** for p *<*0.01, and * for p *<*0.05).

Upon successful purification in detergent micelles (fig. S7A), we determined the specific ATP hydrolysis activity rates as this is a defining catalytic biochemical property of all P-type ATPases. Using *in vitro* malachite green assays, which allow colorimetric quantification of released inorganic phosphate upon ATP hydrolysis, we demonstrated that both GS1 and GS2 exhibit high catalytic turnover rates, despite lacking the MBD. As shown in Fig. 4B, ATPase rate of GS2 (77 *nmol min*^−1^ *mig*^−1^) is comparable to characterized native P_1B-1_-type ATPases (dashed lines) while GS1 (469 *nmol min*^−1^ *mig*^−1^) exceeded those values (*17, 26, 36*). This raised the question of whether the mutations may have compromised the transporter’s ability to autophosphorylate at the conserved DKTGT motif present in the P-domain. To ensure that ATP hydrolysis remained coupled to P-domain phosphorylation, we generated GS1-D349A lacking the conserved aspartate residue which undergoes the required phosphorylation essential to complete the Post-Albers cycle. The absence of any ATPase catalytic activity in GS1-D349A confirmed that ATP hydrolysis in the generated sequence is strictly coupled and dependent on autophosphorylation (Fig. 4B). However, while the ATPase activity in native characterized P_1B-1_-type ATPase is strictly dependent on the presence of Cu(I) substrates, our *in vitro* assay did not show the expected copper dependency. We proposed that this behavior resulted from the *μ*M level copurification of copper with the protein, suggesting a very high affinity of the generated constructs towards Cu(I). This was further supported by the observed reduction but incomplete abolishment of ATP hydrolysis activity upon treatment with up to 50 *μ*M tetrathiomolybdate (TTM), a copper chelator with K_*d*_ of 2.32 × 10^−20^ M (*37*), which effectively competes with Cu(I) availability with the Cu(I)-pumps (Fig. 4B).

Thus, to unambiguously confirm functionality *in vivo*, copper susceptibility and metal quantification assays were performed upon expression of GS1 and GS2 in native and copper-sensitive *E. coli* strains. First, the ability of GS1 and GS2 to efficiently extrude Cu(I) from the cytosol was investigated via *in vivo* intracellular copper quantification by Inductively-Coupled Plasma Mass Spectrometry (ICP-MS) upon cellular growth in media containing increasing copper concentrations. In this assay, we expected functional copper exporters to reduce intracellular copper accumulation. *E. coli* cells that were recombinantly expressing GS1 and GS2 showed reduced copper concentrations compared to cells transformed with a control plasmid not encoding for any Cu(I) P-type ATPase pump (Fig. 4C). In agreement with native-like function, GS1 and GS2’s ability to reduce cellular concentration aligned well with cells overexpressing the functional native *Ec*CopA pump. To further validate the ability of GS1 and GS2 to protect cells from copper toxicity by extrusion, *in vivo* copper metal susceptibility assays were performed in the copper-sensitive *E. coli* ECA464 strain lacking the endogenous bacterial copper efflux and detoxification system genes (*copA, cueO, and cusCFBA*) (*38*). In this assay, protection from copper toxicity was determined by monitoring bacterial growth under increasing copper concentrations to evaluate the protective effect of the expressed transporters. As shown in Fig. 4D (with detailed OD_600_ data from a representative trial in fig. S7B), at a concentration of 6.0 mM copper, cells expressing GS1 (74.6% mean-normalized relative growth) and GS2 (80.8%) demonstrated comparable or exceeding protection ability against copper toxicity than cells expressing the native *Ec*CopA (62.3%). Contrarily, cells transformed with a corresponding empty control vector (14.1%) featured significant growth inhibition due to copper toxicity at higher copper concentrations. Together, experimental *in vitro* and *in vivo* characterization indicate that both generated sequences retain the expected functionality of a P_1B-1_-type ATPase, effectively exporting copper from the cellular cytosol.

## Discussion

Cu(I) transmembrane pumps are complex ATP-dependent multidomain transporters responsible for maintaining essential metal homeostasis in cells in all kingdoms of life. Designing membrane proteins of this level of complexity is significantly challenging because it requires addressing intricate structural dynamics, achieving stability in lipid bilayers, and assessing their functional ability in a native-like environment. Through synergy of machine learning, molecular dynamics, and experimental validation, we designed two artificial transmembrane Cu(I) P-type ATPase pumps (GS1 and GS2), each containing approximately 180 mutations from the referenced WT *Lp*CopA sequence. The LGL framework successfully decoded non-extant novel variants by leveraging preserved co-evolutionary interactions, learned exclusively from the sequence space. The integration of MD and experimentation allowed characterization of essential dynamics and function, providing insights into structural and functional behaviors expected to be preserved in the generated sequences. Results showed that both were capable of adopting the correct fold required for membrane insertion, maintained coupled domain movements, and provided protection against copper toxicity through export across the membrane. The functional efficacy of these artificial transporters marks a major step forward in the field of evolutionary-inspired membrane protein design, offering new possibilities for future advancements.

The interdisciplinary approach presented here can also be applied to other complex, less characterized systems to advance our understanding and engineering of biological systems. Our learning framework integrates an in-depth exploration of sequence space, dynamics, and the biochemical properties of the protein system under consideration. This has significantly narrowed the gap between design predictions and experimental characterization, a trend likely to continue as we deepen our understanding of the functional patterns within protein families as a whole. The potential applications are vast, offering the promise of inspiring numerous additional approaches aimed at designing protein functions in a logical and time-efficient manner.

An important avenue for further investigation includes the analysis of the 32 mutations distinguishing GS1 and GS2 as a potential means of controlling ATP hydrolysis rate. This energy transduction process drives the conformational changes necessary for ion transport. Investigating how these mutations modulate ATP hydrolysis could provide insights into the energy transduction mechanism and overall catalytic efficiency in transmembrane transition metal pumps, and P-type ATPases at large.

## Acknowledgments

The authors thank the Office of Information Technology and the Cyber Infrastructure Research Computing (CIRC) at the University of Texas at Dallas as well as the Texas Advanced Computing Center (TACC) at the University of Texas at Austin for providing HPC resources that have contributed to the research results reported within this paper.

The authors also thank Prof. Dietrich Nies, Martin-Luther University Halle-Wittenberg, Germany, for providing the copper-sensitive *E. Coli* ECA464 strain for this study.

## Author contributions

F.M.O., F.H., G.M., H.T., and F.M. designed research; F.M.O., F.H., and V.V.V. performed research; F.M.O., F.H., V.V.V., G.M., H.T., and F.M. analyzed data; and F.M.O., F.H., G.M., H.T., and F.M. wrote the paper.

## Data and materials availability

The LGL framework used in this study is publicly available at https://github.com/morcoslab/LGL-VAE/. The LGL training dataset, the produced LGL model, a cumulative list of decoded sequences (GS1-GS8) along with MD simulation input files and experimental raw data are uploaded to Zenodo: 10.5281/zenodo.14783470. These files can be accessed through the following link: https://tinyurl.com/4fjwcuhj.

