## Supplementary material for "Generative Landscapes and Dynamics to Design Multidomain Artificial Transmembrane Transporters": Methods and supplementary figures

**List of Supplementary Materials:**

Materials and Methods

Figures S1 to S7

Tables S1 to S2

Supplementary Movies S1 to S2

### Materials and Methods

#### HMMER and HMMSearch for MSA generation

We used the HMMER software package to identify homologous protein and nucleotide sequences. The *HMMSearch* tool generates a multiple sequence alignment (MSA) based on a profile Hidden Markov Model (HMM) that represents patterns associated with a sequence family (*39*). The initial profile HMM is created on the basis of either a sequence or a small MSA consisting of sequences that belong to the family of interest. Databases are searched to identify additional homologous sequences that are appended to the initial MSA and update the statistical model accordingly. After this iterative procedure reaches a threshold, *HMMSearch* stops identifying additional sequences, and a final MSA is obtained.

#### MSA of the $P_{1B}$-type ATPase family

In the CopA system, the metal binding domains (MBDs) have experimentally been demonstrated to only influence the rate of transport but not cargo selectivity and overall translocation abilities, and were thus excluded from the training set (*16*). Furthermore, the inclusion of this domain reduced the resulting number of sequences and diversity in the MSA because it is only associated with certain subgroups (*18*). Using the remaining sequence portions, $P_{1B-1}$ *Legionella pneumophila* CopA (*Lp*CopA) and $P_{1B-2}$ *Shigella sonnei* ZntA (*Ss*ZntA) were aligned to create the initial profile HMM for *HMMSearch*. The master dataset was refined by sub-sampling sequences with below 87 % identity and limiting gaps to 25 continuous positions to reduce redundancy and noise. The resulting MSA comprised 13,553 sequences, with full distribution details provided in fig. S1A. This dataset served as input for training the LGL map shown in Fig. 1B.

#### Direct Coupling Analysis (DCA)

Direct coupling analysis (DCA) is a statistical method that infers direct residue-residue interactions and contacts in the 3D fold of a protein family based on the identification of coevolving residues learned from an MSA (*40*). Each predicted residue-residue interaction is assigned a score called the Direct Information (DI) score, where a higher value indicates more coupled interaction between an amino acid pair. These direct pairs are identified using a Potts model, which quantifies interaction strengths between residues by incorporating both single-site local field parameters $h_i(s_i)$ and pairwise coupling parameters $e_{ij}(s_i, s_j)$ trained on the input MSA (*41*). The parameters $e_{ij}(s_i, s_j)$ quantify the coupling strength between the positions of the residues $i$ and $j$ for all possible pairs of amino acids. Strong coupling pairs represent the likelihood of coevolving relations among residue positions beyond chance. The $h_i(s_i)$ parameters capture the amino acid biases for independent positions, a measure of conservation.

The above parameters can be estimated to derive the Potts model equation below, in which $P(\hat{S})$ represents the probability that the specific sequence $\hat{S}$ belongs to the family represented by the MSA (*40*).

$$P(\hat{S}) = \frac{1}{Z} \exp\left(\sum_i h_i(s_i) + \sum_{i<j} e_{ij}(s_i, s_j)\right) \tag{S1}$$

A Hamiltonian ($H$) equation can be derived from this Potts model, which contains the set of parameters that model the "sequence energy" of the system (*40, 42*):

$$H(\hat{S}) = -\sum_i h_i(s_i) - \sum_{i<j} e_{ij}(s_i, s_j) \tag{S2}$$

In the context of sequences, this Potts Hamiltonian equation measures the fitness of the specific sequence $\hat{S}$ by taking the sum of the $e_{ij}(s_i, s_j)$ and $h_i(s_i)$ parameters determined from the input MSA of the protein family. A greater negative Hamiltonian value predicts relatively higher functional fitness compared to a greater positive value (*42, 43*).

**Variational Auto-Encoder (VAE)**

The Variational Auto-Encoder (VAE) is an unsupervised generative machine learning architecture that compresses high dimensional data into a reduced set of latent variables ($z$). The architecture consists of three parts: 1) encoder, 2) latent space, and 3) decoder. The model is trained upon input data ($S$), which is fed through the encoder network, $q_\phi(z \mid S)$, to learn the features and associated weights that represent the data. Based on these learned parameters, an approximate Gaussian probability distribution is derived to represent the compressed latent space defined by the specified latent variables ($z$). Each latent variable in the latent space is a unique combination of essential features and random noise that represents novel variants of the input dataset. Each set of $z$-variables ($z_0$, $z_1$) from this latent space distribution is fed through the decoder network $p_\theta(\hat{S} \mid z)$, producing the reconstructed data $\hat{S}$.

The overall objective of the architecture is to maximize the evidence lower bound (ELBO) function below (*44*):

$$\mathrm{ELBO}_{\mathrm{VAE}}(\theta, \phi; S) = \mathbb{E}_{q_\phi(z|S)}[\log p_\theta(\hat{S} \mid z)] - \mathrm{KL}[q_\phi(z \mid S) \parallel p(z)] \quad \text{(S3)}$$

In the ELBO, the first term $\mathbb{E}_{q_\phi(z|S)}[\log p_\theta(\hat{S} \mid z)]$ measures the reconstruction quality, while the second term $\mathrm{KL}[q_\phi(z \mid S) \parallel p(z)]$ is the regularization. The Kullback-Leibler (KL) divergence encourages the learned encoder distribution $q_\phi(z \mid S)$ to follow the prior distribution $p(z)$ that is set as a standard Gaussian, thus resulting in the approximate Gaussian probability distribution representative of the latent space.

**Latent Generative Landscape (LGL) framework**

We implemented the Latent Generative Landscape (LGL) architecture, which is an enhanced framework of the VAE that incorporates fitness metrics based on amino acid coevolution, to study protein systems (*14*). The DCA model generates the $e_{ij}(s_i, s_j)$ and $h_i(s_i)$ parameters, while the VAE reconstructs sequences $\hat{S}$ using the same MSA input $S$. Each reconstructed sequence is assigned a fitness score $H(\hat{S})$ based on the DCA parameters, and the final output is the LGL map (Fig. 1A). When visualized in 3D, this map resembles a topographic view, where different colors, valleys, and peaks represent relatively favorable or unfavorable energy for the generated sequences (Fig. 1C).

To visualize the dynamics of fitness across the protein family, a 500 x 500 coordinate grid LGL is produced using a curated MSA as input as demonstrated in Fig. 1A *right*. Each pixel in the map represents a reconstructed sequence $\hat{S}$ from the VAE. Each reconstructed sequence is scored based on its Hamiltonian value $H(\hat{S})$ with respect to the family and is color-coded accordingly (*40, 42*). The dark native blue valleys represent low-energy, favorable configurations while the bright green peaks indicate sequences less explored by the extant dataset.

Overlaying native input sequences onto the generated LGL map provides deeper insights into phylogenetic relationships and the distribution of features across the latent space (*2, 13, 14*). Fig. 1B presents the LGL of the $P_{1B}$-type ATPase family, with colored symbols representing labeled sequences from the input training MSA, categorized by their corresponding subgroups. Sequences within valleys or clusters share common characteristics, allowing a chance to explore underlying patterns previously unidentified within a protein family (*3*). Novel reconstructed sequences $\hat{S}$ can be decoded based on fitness score and shared characteristics with overlapping native family sequences $S$.

**Molecular dynamics system preparation**

The starting structures (*Lp*CopA, GS1-GS8) were generated by homology modeling with the E2$P_i$ template structure (PDB: 3RFU) using SWISS-MODEL (*45, 46*). Afterward, protonation states at pH 7.4 were assigned using the H++ online server (*47–49*). Additionally, protein orientations within the lipid bilayer were determined using the Positioning of Proteins in Membranes (PPM2.0) algorithm (*29*). Subsequently, each system was prepared for MD simulations using the Amber20 suite (*50*). This involved using the *tleap* program to cap the termini accordingly, while *packmol-memgen* embedded the proteins in a lipid bilayer with a composition of six 1-palmitoyl-2-oleoyl-sn-glycero-3-phosphoethanolamine (POPE), three 1-palmitoyl-2-oleoyl-sn-glycero-3-phosphocholine (POPC), two 1-Palmitoyl-2-oleoyl-sn-glycero-3-phosphoglycero (POPG), and one 1,3-bis(1-oleoyl-2-palmitoyl-sn-glycero-3-phospho)-sn-glycerol (OPOPCL). The systems were solvated with water and $Na^+$ ions were added to neutralize the negatively charged phospholipid heads (*50–52*). Then, *tleap* was used again to introduce a 150 mM NaCl concentration atop the neutralizing ions and to generate the parameter and coordinate files. The force fields used included ff14SB, lipid17, lipid17$ext$, and TIP3P for water (*53–56*).

**Molecular dynamics simulation protocol**

MD simulations were conducted using the GPU-accelerated *pmemd* program within the Amber20 suite (*50*). Initial energy minimization was carried out in four stages, each consisting of 5000 steps of steepest descent followed by 5000 steps of conjugate-gradient minimization, progressively reducing the amount of restraints. Restraints of 20 kcal/mol/$\AA^2$ were applied as follows: first, on all atoms except lipids; then to protein and water atoms; next, exclusively to protein atoms; and, finally, with no restraints. The systems were heated in two phases: from 0 to 100K over 6 ns, and from 100K to 303K over 12 ns, using a Langevin thermostat while maintaining the lipid bilayer and protein under soft restraints (5 kcal/mol/$\AA^2$) (*57*). After heating, a membrane equilibration protocol was applied with ten steps of 500 ps to stabilize the periodic boundary conditions. Subsequently, production of MD simulations was run in trials of 400 ns, each under the NPT ensemble with a 2 fs time-step, Langevin dynamics, a 12 Å cutoff for nonbonded interactions, and the SHAKE algorithm (*58*). Data analysis was performed using the *cpptraj* module in Amber20 and Tcl script for VMD (*50, 59, 60*).

**A-domain conformational analysis**

The transmembrane helices in each trajectory were aligned with those in the initial structure of the E2$P_i$ state of *Lp*CopA (PDB: 3RFU) to ensure consistency and accurate measurement. Two metrics were computed using an in-house Tcl script for VMD to capture the movement of the A domain: the tilt angle and the displacement distance denoted as Δdistance. The tilt angle was calculated by taking the cross product of two orthogonal vectors, which are highlighted by the pink dashed lines in Fig. 3B. The first vector, representing a stable horizontal reference perpendicular to the membrane normal vector, was defined along the MB' helix kink at the lipid bilayer-solvent interface. The second vector extended from one of the basal $\alpha$-helices to the center of mass of the A-domain's $\beta$-sheets, providing a direct measure of domain tilt relative to the membrane plane. The second metric, Δdistance, was defined as the distance between the center of mass of the A-domain in each frame and its center of mass in the E1 state (PDB: 7R0I), depicted as the blue solid line in Fig. 3B. The Δdistance allowed us to evaluate any translational movement of the domain toward the E1 state. *Lp*CopA and GS1-GS8 results for each trial are presented in fig. S5C and S6C. To streamline interpretation, these metrics were combined into an *A-domain movement score* as follows:

1. Frame vector construction: the frames' angle and displacement distance scalar values ($\theta$ and Δd) were combined to create 2D vectors.

$$\text{Frame vector} = \mathbf{F_j} = (F_{j\theta}, F_{j\Delta d}) \tag{S4}$$

2. Frame Euclidean distance: for each frame, the Euclidean distance between the frame vector and the target vector (E1) was calculated as:

$$\text{Frame Euclidean distance} = ||\mathbf{E1} - \mathbf{F_j}|| = \sqrt{(E1_\theta - F_{j\theta})^2 + (E1_{\Delta d} - F_{j\Delta d})^2} \tag{S5}$$

3. Target Euclidean distance: the Euclidean distance between the reference vector (E2$P_i$) and the target vector (E1) was calculated as shown:

$$\text{Target Euclidean distance} = ||\mathbf{E1} - \mathbf{E2P_i}|| = \sqrt{(E1_\theta - E2P_{i\theta})^2 + (E1_{\Delta d} - E2P_{i\Delta d})^2} \tag{S6}$$

4. Movement score calculation: the frame Euclidean distance was compared to the target Euclidean distance to calculate the *A-domain movement score* for each frame:

$$\textit{A-domain movement score} = \left(1 - \frac{\|\mathbf{E1} - \mathbf{F_j}\|}{\|\mathbf{E1} - \mathbf{E2P_i}\|}\right) \times 100 \qquad \text{(S7)}$$

The resulting probability density of the *A-domain movement scores* provides a comprehensive measure of A-domain conformational change (Fig 3D and fig. S5B). Positive values signify movement toward the E1 state and are labeled as "progress," while negative values indicate movement away from the E1 state and are labeled as "deviation." Additionally, table S1 summarizes the maximum coupled A-domain progress achieved in each sequence trial.

**Transmembrane helix rearrangement analysis**

A-domain rearrangement is functionally meaningful only if it induces conformational changes in the transmembrane (TM) domain. In $P_{1B-1}$-type ATPases, these conformational changes involve the relative movement of two distinct blocks, each containing four transmembrane $\alpha$-helices (MA, MB, M1, M2) and (M3, M4, M5, M6). To quantify this movement, we performed both qualitative and quantitative analyses. For qualitative analysis, we visualized the superposition of the two TM blocks (colored green and purple) at various stages of the A-domain progress, relative to the reference starting structure of *Lp*CopA (PDB: 3RFU), depicted in lighter shades of the same colors in Fig. 3E. This visualization provided an intuitive representation of the directional shifts in TM blocks over time. For quantitative analysis, we utilized the *pyDDM* tool to compute a distance difference matrix (DDM), enabling detailed measurement of both intra-block and inter-block changes across trajectory frames relative to the reference structure (*61*). This matrix allowed us to detect inter-block rearrangements with high precision with table S2 showcasing the maximum coupled inter-block DDM values featured in each sequence trial. The combination of quantitative distance analysis and qualitative visualization provided a comprehensive picture of TM block movement and confirmed whether the block shifts remained aligned with the *A-domain movement score*. Additionally, CAVER 3.0 analysis was also performed to monitor binding site accessibility on both the intracellular and extracellular sides adding another layer of verification for TM helix rearrangement. According to the Post-Albers cycle, simulations exhibiting large A-domain transitions are expected to show transmembrane rearrangements that occlude the release pathway while subsequently exposing the uptake pathway, as shown in Fig. 3E.

#### Molecular dynamics results of additional decoded generated sequences

To better understand the impact of the Hamiltonian metric on the generated protein dynamics, we conducted a similar analysis on additional sequences from across the *Lp*CopA area. They were categorized as GS3 to GS8 and their locations on the LGL map and Hamiltonian assigned value can be found in fig. S5, A and D.

Regarding their A-domain's ability to undergo conformational changes, several additional decoded sequences did not exhibit large dynamics. First, we observed that various sequences (GS5-GS7) no longer exhibit the characteristic bimodal distribution of the WT in the probability distribution of *A-domain movement scores* presented in fig. S5B. Furthermore, when considering coupled motions between the soluble domain and the TM blocks, half of the additional selected sequences underperform as shown in table S1. Particularly GS4-GS6 do not reach a maximum coupled *A-domain movement score* of 50.00 (half progress). Even when selecting a pliable *A-domain movement score* threshold of 45 based on WT simulation results as well as to accommodate small simulation deviations, the trials of GS4-GS6 were deemed not dynamic enough.

In studying TM helix rearrangement, we found that most of these sequences lacked clear inter-block differentiation (fig. S5D). In several cases, a fraction of the block exhibited significant movement, while the block as a whole did not shift, suggesting a weak intra-block correlation. This observation was quantified by measuring the larger standard deviation between rows and columns per inter-block DDM. Moreover, when contemplating coupled rearrangements between the A-domain and TM domain, approximately half of the trials exhibited underachieving inter-block DDM scores or intra-block correlations. When selecting soft thresholds based on WT simulation results and slight simulation fluctuations, an inter-block DDM score of 1.80 $\mathring{A}$ and inter-block DDM standard deviation smaller than 0.75 $\mathring{A}$, only GS3 and GS4 displayed consistently significant coupled inter-block DDM scores.

The additional generated sequences (GS3-GS8) combined results of coupled *A-domain movement score* and TM inter-block rearrangement analyses demonstrated subpar inter-domain communication. The same chosen thresholds showcased that half of the WT *Lp*CopA, GS1, and GS2 simulation trials met the requirements whereas the remaining generated sequences did not. These results could indicate that GS3-GS8 decoded sequences could have slow or inefficient coupled dynamics, while GS1 and GS2 larger maximum coupled values resembling WT *Lp*CopA dynamics suggested faster and more coupled dynamics (tables S1 and S2). Thus, to optimize experimental resources, we prioritized GS1 and GS2 for experimental characterization.

**Protein expression of selected generated sequences**

The codon optimized synthetic DNA encoding GS1 and GS2 sequences (Genscript Inc.) was cloned into a pET-52b(+) vector with an N-terminal STREP-tag II and a C-terminal His-Tag. The constructs were then transformed into *E. coli* BL21-Gold(DE3) competent cells for recombinant protein expression. Transformed cells were grown aerobically overnight in Terrific Broth (TB) media supplemented with 50 $\mu$g/mL ampicillin. The overnight cultures were subsequently inoculated into fresh TB media containing ampicillin (50 $\mu$g/mL) and grown aerobically at 37°C with shaking (240 rpm) until they reached $OD_{600}$ 2.0. Protein expression was induced by the addition of 0.5 mM IPTG and cells were incubated at 30°C for an additional 6 hours. Cells were harvested by centrifugation at 4,000 x*g* for 20 minutes at 4°C.

Harvested cells were resuspended in lysis buffer (20 mM Tris-HCl pH 8, 150 mM NaCl, 5mM $MgCl_2$, 30 $\mu$g/mL DNaseI from bovine pancreas, and EDTA-free protease inhibitor cocktail (Thermo Scientific)), and lysed using an ice-cooled microfluidizer at 20,000 psi for four cycles (Microfluidics M-110P). The lysate was subjected to two centrifugation steps: first at 20,000 x*g* for 20 minutes at 4°C to remove cell debris and subsequently at 180,000 x*g* for 1 hour at 4°C to pellet the cell membranes. The membrane pellet was resuspended in buffer (20 mM Tris-HCl pH 8, 500 mM NaCl, 10% (w/v) glycerol, and EDTA-free protease inhibitor cocktail (Thermo Scientific)), flash-frozen in liquid nitrogen, and stored at -80°C until purification. Also, to confirm protein expression, the resuspended membranes and supernatant from the last centrifugation were subjected to SDS-PAGE and Western Blot. Samples containing approximately 15 $\mu$g of total protein were subjected to SDS-PAGE (4-15%). Consequently, protein bands were turbo-transfered onto a nitrocellulose membrane, which was immediately blocked using 3% non-fat milk in TBST buffer (Tris Buffered Saline buffer with a 0.1% concentration of Tween 20) at room temperature for 1 hour. Then, the primary antibody against STREP-tag II was added in 3% milk in TBST onto the membrane and incubated overnight at 4°C. Afterward, the membrane was rinsed with TBST buffer, and the alkaline-phosphatase conjugated secondary anti-mouse IgG antibody in 3% milk in TBST buffer was poured onto the membranes for 2 hours at room temperature. Lastly, after rinsing the membranes with TBST the bands were developed using nitroblue tetrazolium (NBT) and 5-bromo-4-chloro-3-indolyl phosphate (BCIP) in Tris buffer.

#### Protein purification of selected generated sequences

Purification followed an established protocol for *Ec*CopA (*17*). Briefly, membrane proteins were extracted by incubating the membrane suspension with 1% (w/v) n-dodecyl-$\beta$-d-maltoside (DDM) detergent in buffer (20 mM Tris-HCl pH 8, 500 mM NaCl, 25 mM imidazole, 5 mM $\beta$-mercaptoethanol and EDTA-free protease inhibitor cocktail) for 1 hour at 4°C, followed by ultracentrifugation at 180,000 x*g* for 30 minutes at 4°C to remove residual membranes. The detergent-solubilized supernatant was loaded onto a 5 mL HisTrap affinity column (Cytiva) at 0.5 mL/min flow rate in binding buffer (20 mM Tris-HCl pH 8, 500 mM NaCl, 25 mM imidazole, 1 mM DTT, and 0.05 % (w/v) DDM). Protein elution was achieved with elution buffer (20 mM Tris-HCl pH 8, 500 mM NaCl, 400 mM imidazole, 1 mM DTT, and 0.05 % (w/v) DDM). The eluted protein was loaded onto a HiPrep 26/10 desalting column (Cytiva) equilibrated with 20 mM MOPS/HCl pH 7, 500 mM NaCl, 1mM EDTA, 1 mM DTT, and 0.05 % (w/v) DDM to remove imidazole and exchange the buffer. The protein was concentrated using a 100 kDa MWCO spin concentrator to 2 mg/mL and afterward loaded onto a Superdex S200 10/300 size exclusion chromatography column (Cytiva) for isolation of monodisperse protein using sizing buffer (20 mM MOPS/HCl pH 7, 500 mM NaCl, 1 mM DTT, and 0.05 % (w/v) DDM). SDS-PAGE (4-15%) confirmed purity of the protein, and concentration was determined by $Abs_{280}$ (GS1 MW=71.6 KDa and $\varepsilon_{280}$=58,440 $M^{-1}cm^{-1}$, GS2 MW=70.4 KDa and $\varepsilon_{280}$=56,950 $M^{-1}cm^{-1}$). The purified protein samples were flash-frozen in liquid nitrogen and stored at -80°C until use.

#### *In vitro* ATPase assays

ATPase hydrolysis assays were conducted using a Malachite Green Phosphate Assay Kit (Sigma Millipore). Protein samples were diluted to 0.01 mg/mL in a buffer containing 20 mM MOPS/HCl pH 7, 500 mM NaCl, 1 mM DTT, and 0.05 % (w/v) DDM. All solutions used in the assay were prepared with Chelex-treated Mili-Q water to minimize trace metals contamination. Each assay sample (200 $\mu$L), was obtained by mixing 172 $\mu$L of 0.01 mg/mL protein, 16 $\mu$L of buffer, 2 $\mu$L of a 100X $CuCl_2$ stock, 2 $\mu$L of TTM (to a final concentration of 50 $\mu$M) or DMSO, 4 $\mu$L of cysteine (final concentration 2 mM), 2 $\mu$L of $MgCl_2$ (final concentration 10 mM), and 2 $\mu$L of ATP (final concentration 1mM) to initiate the reaction. The reaction was conducted at 37°C under continuous agitation for 10 minutes. Following incubation, 100 $\mu$L of the kit's Working Reagents (Molybdate and Malachite Green) were added to each well to enable color development, proportional to inorganic phosphate ($P_i$) produced during ATP hydrolysis. After transferring samples to a 96-well plate, absorbance at 620 nm was measured using a Tecan Spark 20 M plate reader. To quantify $P_i$, a calibration curve was generated using the $P_i$ standard provided in the assay kit. Eight standards ranging from 0 to 40 $\mu$M were prepared in 200 $\mu$L volumes. ATPase activity was calculated and reported as nmol of $P_i$ per mg of protein per minute.

#### ***In vivo*** **intracellular copper quantification assays**

Overnight cultures of *E. coli* BL21-Gold(DE3) cells transformed with expression plasmids encoding GS1, GS2, *Ec*CopA, or control (pET-52b(+)), were inoculated into 35 mL of Terrific Broth (TB) media supplemented with 50 $\mu$g/mL ampicillin and grown aerobically until reaching an $OD_{600}$ of 2.0 at 37°C. Protein expression was induced by adding 0.5 mM IPTG, and the cultures were incubated for 6 hours at 30°C. Subsequently, 10% (v/v) of the cultures was transferred into pre-weighted tubes containing 5 mL of TB media with ampicillin (50 $\mu$g/mL) and $CuCl_2$ at final concentrations ranging from 0 mM to 2 mM, with each condition performed in triplicate. These cultures were then grown aerobically under continuous shaking for 10 hours overnight at 30°C. The cells were washed to remove excess extracellular copper by pelleting via centrifugation at 4,000 x*g* for 20 minutes at 4°C, resuspending in 5 mL of copper-free TB media, and repeating centrifugation. The resulting cell pellets were dried overnight in a vacuum chamber, and tube weights were recorded to calculate the cell mass. The pellets were then digested in 50% nitric acid (v/v) and incubated at 80°C overnight. Samples were subsequently diluted to a 3% nitric acid (v/v) before copper quantification via Inductively-Coupled Plasma Mass Spectrometry (ICP-MS) measurements, with intracellular copper levels reported as $\mu$g of Cu per mg of cell mass. Metal content was analyzed with an Agilent 7900 ICP mass spectrometer connected to a CETACASX-500 auto-sampler for sample injection. Statistical significance was evaluated using an unpaired *t*-test, on two independent biological replicates.

***In vivo* copper susceptibility growth assays**

To better elucidate the impact of GS1 and GS2 expression in copper extrusion and thereby cellular protection from copper toxicity, copper sensitive *E. coli* strain ECA464 was selected for copper susceptibility toxicity assays (*38*). The copper sensitive strain was subjected to a lysogenization with $\lambda$DE3 phage following the manufacturer's user protocol (Novagen cat. 69734). Furthermore, ECA464 $\lambda$DE3 lysogens were made competent prior to plasmid transformation. The expression of recombinant proteins in this strain was confirmed prior to the assay via western blot using the previously described protocol (Fig. 4A). Thereafter, overnight cultures of transformed cells were inoculated into fresh TB media supplemented with 50 $\mu$g/mL ampicillin and grown at 37°C to an $OD_{600}$ of 2.0. Protein expression was induced by adding 0.5 mM IPTG, and cultures were incubated for an additional 5 hours at 30°C. To monitor the effect of the recombinantly expressed protein, a 96-well plate setup was employed to track $OD_{600}$ cell growth at increasing copper concentrations. Each well was supplemented with TB media containing ampicillin (50 $\mu$g/mL), IPTG (0.5 mM) and varying concentrations of $CuCl_2$ (0 to 6 mM). Additionally, each well was filled with its corresponding cell culture at a 10% (v/v) dilution. Each construct was tested in triplicate for each copper concentration. $OD_{600}$ was measured in 10-minute intervals over 16 hours at 37°C to monitor cell growth using a Tecan Spark 20 M plate reader. See fig. S7B for $OD_{600}$ growth curves. The data was further analyzed by calculating the mean-normalized relative $OD_{600}$ growth for each concentration, using the highest absorbance value recorded at the stationary phase as the reference point (defined as 100% growth). Four biological replicates were performed, and statistical significance was assessed with an unpaired *t*-test.

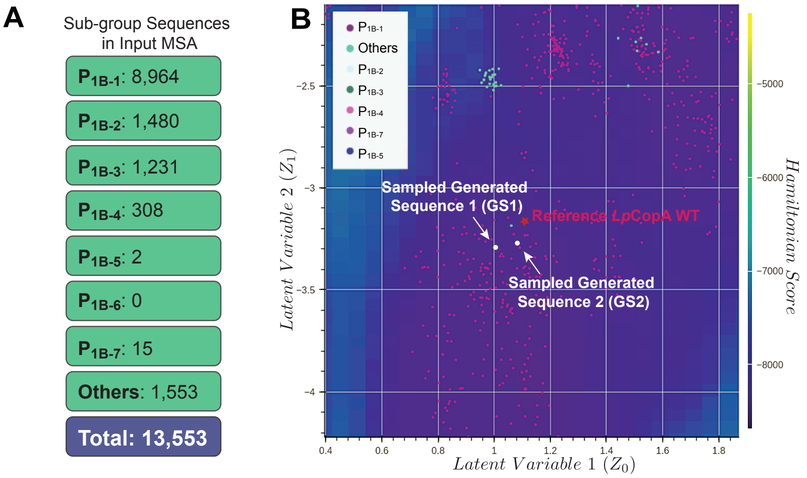


**Figure S1**: **LGL training input MSA summary and native sequences placement near GS1 and GS2.** (**A**) Sequence distribution of the $P_{1B}$-ATPase family used to train the LGL model. (**B**) A 2D close-up of the reference *Lp*CopA WT labeled with a red star. The two selected decoded sequence (GS1 and GS2) locations are labeled with white circles. The nearby colored dots represent other training sequences showing no overlap with the decoded sequences.

```
            MA                                       MB
            1            10              20              30              40              50              60
GS1         YLDMTRRFWIGLALTLPVFVLEMGGHGLHHLIDPQLSNWIQLALATPVVLWAGWPFFVRG
GS2         .VDMTRRFWIGLALTLPVFVLEMGGHGLHHLIGPQLSNWIQLALATPVVLWAGWPFFVRG
LpCopA_WT   YLDMRRRFWIALMLTIPVVILEMGGHGLKHFISGNGSSWIQLLLATPVVLWGGWPFFKRG
```

```
                           M1                                       M2
                        70              80              90             100             110             120
GS1         WASVRTRNLNMFTLIALGTGVAWLYSVVATLAPGLFPPAFRDHDGAVAVYFEAAAVITVL
GS2         WQSLVTRNLNMFTLIALGTGVAWLYSVVATLAPGLFPPAFRGHDGAVAVYFEAAAVITVL
LpCopA_WT   WQSLKTGQLNMFTLIAMGIGVAWIYSMVAVLWPGVFPHAFRSQEGVVAVYFEAAAVITTL
```

```
                           A-Domain
                       130             140             150             160             170             180
GS1         VLLGQVLELRARERTSGAIRALLDLAPKTARRIGADGSEEEVALDQVQVGDRLRVRPGEK
GS2         VLLGQVLELRARERTGGAIRALLDLAPKTARRIGADGSEEEVPLDQVVVGDRLRVRPGEK
LpCopA_WT   VLLGQVLELKAREQTGSAIRALLKLVPESAHRIKEDGSEEEVSLDNVAVGDLLRVRPGEK
```

```
                       190             200             210             220             230             240
GS1         VPVDGEVLEGRSSVDESMVTGESMPVTKEVGDKVIGGTINQTGSFVMRAEKVGRDTMLSR
GS2         VPVDGEVLEGRSSVDESMVTGESMPVTKEAGDKVIGGTINQTGSFVMRAEKVGADTMLSQ
LpCopA_WT   IPVDGEVQEGRSFVDESMVTGEPIPVAKEASAKVIGATINQTGSFVMKALHVGSDTMLAR
```

```
                                M3                                          M4
                       250             260             270             280             290             300
GS1         IVQMVAQAQRSRAPIQRLADQVSGWFVPAVIAVALLAFAAWALFGPEPRFSYALIAAVSV
GS2         IVQMVAEAQRSRAPIQRLADQVSGWFVPAVIAVALLAFAAWAIFGPEPAFSYALIAAVSV
LpCopA_WT   IVQMVSDAQRSRAPIQRLADTVSGWFVPAVILVAVLSFIVWALLGPQPALSYGLIAAVSV
```

```
                                                       P-Domain
                       310             320             330             340             350             360
GS1         LIIACPCALGLATPMSIMVGVGRGAQAGVLIKNAEALERMEKVDTLVVDKTGTLTEGKPK
GS2         LIIACPCALGLATPMSIMVGVGRGAQAGVLIKNAEALERMEKVDTLVVDKTGTLTEGKPK
LpCopA_WT   LIIACPCALGLATPMSIMVGVGKGAQSGVLIKNAEALERMEKVNTLVVDKTGTLTEGHPK
```

```
            N-Domain
                       370             380             390             400             410             420
GS1         VTAVVPAAGFDEAELLRLAASLERASEHPLAAAIVAAAEERGLTLAEVEDFDSPTGKGVT
GS2         VTAVVPAAGFAEAELLRLAASLERGSEHPLAAAIVAAAEERGLTLAEVEDFDSPTGKGVT
LpCopA_WT   LTRI.VTDDFVEDNALALAAALEHQSEHPLANAIVHAAKEKGLSLGSVEAFEAPTGKGVV
```

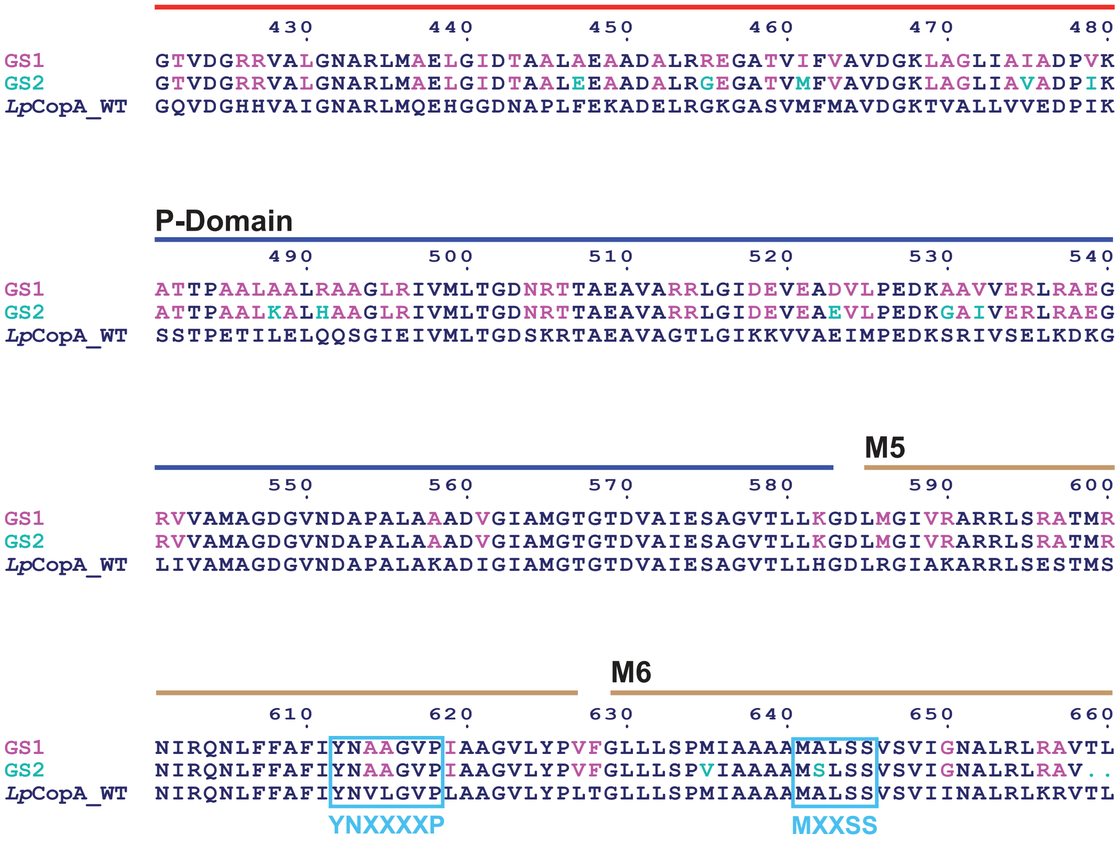


**Figure S2**: **Sequence alignment of GS1, GS2, and WT *Lp*CopA**. The color scheme for mutations corresponds to that of Fig. 2C: magenta indicates mutations from *Lp*CopA, while teal highlights the 32 differences between GS1 and GS2. Additionally, domain boundaries and key motifs are labeled for clarity. The LGL decoded sequences were used as alignments against *Lp*CopA without modification, and the figure was generated using ESPript3 (*62*).

```
MA  MB
                  1         10        20        30        40        50        60
XtCopA|A0A6I8R0A5 IKQWRNSFLFSLLFGIPVIILMIYMLAANKDHHNTMVLDRNIVPGLSIINLVFFILCTFV
HumanATP7B|P35670 IKQWKKSFLCSLVFGIPVMALMIYMLIPSNEPHQSMVLDHNIIPGLSILNLIFFILCTFV
AfCopA|O29777     ..............................LA.......HFIS..LPYEDFVQLLIALPA
EcCopA|Q59385     ......QAIVALAVGIPVMVWGMI...GDNMM.......V.TA..DNRSLWLVILITLAV
LpCopA_WT|Q5ZWR1  YLDMRRRFWIALMLTIPVVILEMG...GHGLK.......HFIS..GNGSSWIQLLLATPV
GS1               YLDMTRRFWIGLALTLPVFVLEMG...GHGLH.......HLID..PQLSNWIQLALATPV
GS2               .VDMTRRFWIGLALTLPVFVLEMG...GHGLH.......HLIG..PQLSNWIQLALATPV

M1  M2
                            70        80        90       100       110
XtCopA|A0A6I8R0A5 QTLGGRYFYVQAYKSLKHKATNMDVLIVLATTIAYIYSVVILTVAMV.....EKADKSPE
HumanATP7B|P35670 QLLGGWYFYVQAYKSLRHRSANMDVLIVLATSIAYVYSLVILVVAVA.....EKAERSPV
AfCopA|O29777     IFYSGSSIFKAAFSALRRRTLNMDVMYSMGVGAAFLASVLS..TAGVLPREYS.......
EcCopA|Q59385     MVFAGGHFYRSAWKSLLNGAATMDTLVALGTGVAWLYSMSVNLWPQWFPMEAR......H
LpCopA_WT|Q5ZWR1  VLWGGWPFFKRGWQSLKTGQLNMFTLIAMGIGVAWIYSMVAVLWPGVFPHAFRSQEGVVA
GS1               VLWAGWPFFVRGWASVRTRNLNMFTLIALGTGVAWLYSVVATLAPGLFPPAFRDHDGAVA
GS2               VLWAGWPFFVRGWQSLVTRNLNMFTLIALGTGVAWLYSVVATLAPGLFPPAFRGHDGAVA

A-Domain
                    120       130       140       150       160       170
XtCopA|A0A6I8R0A5 TFFDTPPMLFMFIALGRWLEHIAKSKTSEALAKLISLQATEAAVVTFGANQIILREEQVA
HumanATP7B|P35670 TFFDTPPMLFVFIALGRWLEHLAKSKTSEALAKLMSLQATEATVVTLGEDNLIIREEQVP
AfCopA|O29777     .FYETSVLLLAFLLLGRTLEARAKSRTGEAIKKLVGLQAKTAVVIR.DG.....KEIAVP
EcCopA|Q59385     LYYEASAMIIGLINLGHMLEARARQRSSKALEKLLDLTPPTARLVTDEG.....E.KSVP
LpCopA_WT|Q5ZWR1  VYFEAAAVITTLVLLGQVLELKAREQTGSAIRALLKLVPESAHRIKEDG.....SEEEVS
GS1               VYFEAAAVITVLVLLGQVLELRARERTSGAIRALLDLAPKTARRIGADG.....SEEEVA
GS2               VYFEAAAVITVLVLLGQVLELRARERTGGAIRALLDLAPKTARRIGADG.....SEEEVP

                    180       190       200       210       220       230
XtCopA|A0A6I8R0A5 VELVQRGDIVKVVPGGKFPVDGKVIEGTSMADESLITGEPMPVRKKPGSMVIAGSINAHG
HumanATP7B|P35670 MELVQRGDIVKVVPGGKFPVDGKVLEGNTMADESLITGEAMPVTKKPGSTVIAGSINAHG
AfCopA|O29777     VEEVAVGDIVIVRPGEKIPVDGVVVEGESYVDESMISGEPVPVLKSKGDEVFGATINNTG
EcCopA|Q59385     LAEVQPGMLLRLTTGDRVPVDGEITQGEAWLDEAMLTGEPIPQQKGEGDSVHAGTVVQDG
LpCopA_WT|Q5ZWR1  LDNVAVGDLLRVRPGEKIPVDGEVQEGRSFVDESMVTGEPIPVAKEASAKVIGATINQTG
GS1               LDQVQVGDRLRVRPGEKVPVDGEVLEGRSSVDESMVTGESMPVTKEVGDKVIGGTINQTG
GS2               LDQVVVGDRLRVRPGEKVPVDGEVLEGRSSVDESMVTGESMPVTKEAGDKVIGGTINQTG
                                                       T/SGE

M3
                    240       250       260       270       280       290
XtCopA|A0A6I8R0A5 TVLVEATHVGSETTLAQIVKLVEEAQMSKAPIQQLADKISGYFVPFIIIISVVTLVTWII
HumanATP7B|P35670 SVLIKATHVGNDTTLAQIVKLVEEAQMSKAPIQQLADRFSGYFVPFIIIMSTLTLVVWIV
AfCopA|O29777     VLKIRATRVGGETLLAQIVKLVEDAMGSKPPIQRLADKVVAYFIPTVLLVAISAFIYWYF
EcCopA|Q59385     SVLFRASAVGSHTTLSRIIRMVRQAQSSKPEIGQLADKISAVFVPVVVVIALVSAAIWYF
LpCopA_WT|Q5ZWR1  SFVMKALHVGSDTMLARIVQMVSDAQRSRAPIQRLADTVSGWFVPAVILVAVLSFIVWAL
GS1               SFVMRAEKVGRDTMLSRIVQMVAQAQRSRAPIQRLADQVSGWFVPAVIAVALLAFAAWAL
GS2               SFVMRAEKVGADTMLSQIVQMVAEAQRSRAPIQRLADQVSGWFVPAVIAVALLAFAAWAI
```

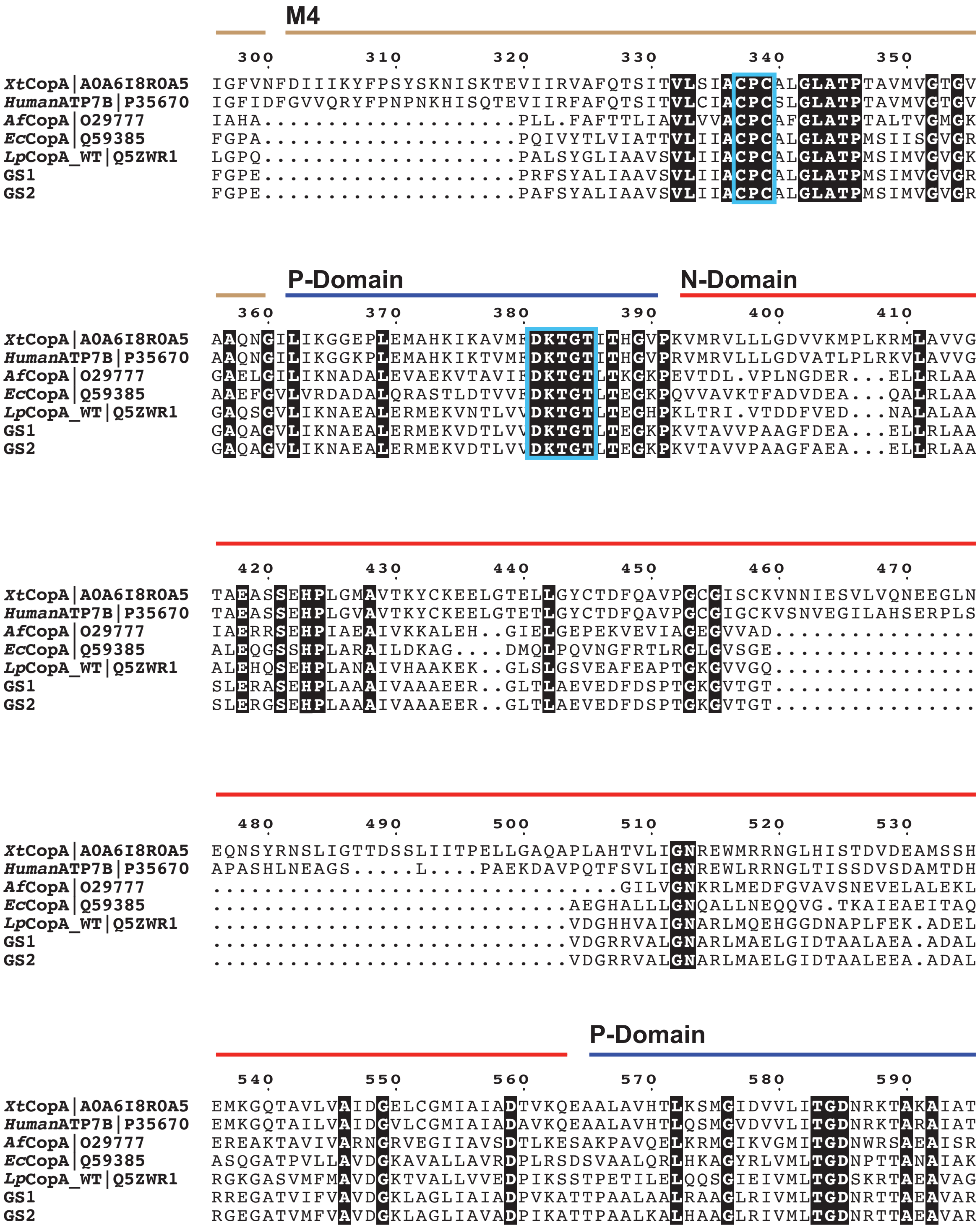
M4
300 310 320 330 340 350
XtCopA|A0A6I8R0A5 IGFVNFDIIIKYFPSYSKNISKTEVIIRVAFQTSITVLSIACPCALGLATPTAVMVGTGV
HumanATP7B|P35670 IGFIDFGVVQRYFPNPNKHISQTEVIIRFAFQTSITVLCIACPCSLGLATPTAVMVGTGV
AfCopA|O29777 IAHA....................PLL.FAFTTLIAVLVVACPCAFGLATPTALTVGMGK
EcCopA|Q59385 FGPA....................PQIVYTLVIATTVLIIACPCALGLATPMSIISGVGR
LpCopA_WT|Q5ZWR1 LGPQ....................PALSYGLIAAVSVLIIACPCALGLATPMSIMVGVGK
GS1 FGPE....................PRFSYALIAAVSVLIIACPCALGLATPMSIMVGVGR
GS2 FGPE....................PAFSYALIAAVSVLIIACPCALGLATPMSIMVGVGR
P-Domain
N-Domain
360 370 380 390 400 410
XtCopA|A0A6I8R0A5 AAQNGILIKGGEPLEMAHKIKAVMFDKTGTITHGVPKVMRVLLLGDVVKMPLKRMLAVVG
HumanATP7B|P35670 AAQNGILIKGGKPLEMAHKIKTVMFDKTGTITHGVPRVMRVLLLGDVATLPLRKVLAVVG
AfCopA|O29777 GAELGILIKNADALEVAEKVTAVIFDKTGTLTKGKPEVTDL.VPLNGDER...ELLRLAA
EcCopA|Q59385 AAEFGVLVRDADALQRASTLDTVVFDKTGTLTEGKPQVVAVKTFADVDEA...QALRLAA
LpCopA_WT|Q5ZWR1 GAQSGVLIKNAEALERMEKVNTLVVDKTGTLTEGHPKLTRI.VTDDFVED...NALALAA
GS1 GAQAGVLIKNAEALERMEKVDTLVVDKTGTLTEGKPKVTAVVPAAGFDEA...ELLRLAA
GS2 GAQAGVLIKNAEALERMEKVDTLVVDKTGTLTEGKPKVTAVVPAAGFAEA...ELLRLAA
420 430 440 450 460 470
XtCopA|A0A6I8R0A5 TAEASSEHPLGMAVTKYCKEELGTELLGYCTDFQAVPGCGISCKVNNIESVLVQNEEGLN
HumanATP7B|P35670 TAEASSEHPLGVAVTKYCKEELGTETLGYCTDFQAVPGCGIGCKVSNVEGILAHSERPLS
AfCopA|O29777 IAERRSEHPIAEAIVKKALEH..GIELGEPEKVEVIAGEGVVAD................
EcCopA|Q59385 ALEQGSSHPLARAILDKAG....DMQLPQVNGFRTLRGLGVSGE................
LpCopA_WT|Q5ZWR1 ALEHQSEHPLANAIVHAAKEK..GLSLGSVEAFEAPTGKGVVGQ................
GS1 SLERASEHPLAAAIVAAAEER..GLTLAEVEDFDSPTGKGVTGT................
GS2 SLERGSEHPLAAAIVAAAEER..GLTLAEVEDFDSPTGKGVTGT................
480 490 500 510 520 530
XtCopA|A0A6I8R0A5 EQNSYRNSLIGTTDSSLIITPELLGAQAPLAHTVLIGNREWMRRNGLHISTDVDEAMSSH
HumanATP7B|P35670 APASHLNEAGS.....L....PAEKDAVPQTFSVLIGNREWLRRNGLTISSDVSDAMTDH
AfCopA|O29777 ................................GILVGNKRLMEDFGVAVSNEVELALEKL
EcCopA|Q59385 ............................AEGHALLLGNQALLNEQQVG.TKAIEAEITAQ
LpCopA_WT|Q5ZWR1 ............................VDGHHVAIGNARLMQEHGGDNAPLFEK.ADEL
GS1 ............................VDGRRVALGNARLMAELGIDTAALAEA.ADAL
GS2 ............................VDGRRVALGNARLMAELGIDTAALEEA.ADAL
P-Domain
540 550 560 570 580 590
XtCopA|A0A6I8R0A5 EMKGQTAVLVAIDGELCGMIAIADTVKQEAALAVHTLKSMGIDVVLITGDNRKTAKAIAT
HumanATP7B|P35670 EMKGQTAILVAIDGVLCGMIAIADAVKQEAALAVHTLQSMGVDVVLITGDNRKTARAIAT
AfCopA|O29777 EREAKTAVIVARNGRVEGIIAVSDTLKESAKPAVQELKRMGIKVGMITGDNWRSAEAISR
EcCopA|Q59385 ASQGATPVLLAVDGKAVALLAVRDPLRSDSVAALQRLHKAGYRLVMLTGDNPTTANAIAK
LpCopA_WT|Q5ZWR1 RGKGASVMFMAVDGKTVALLVVEDPIKSSTPETILELQQSGIEIVMLTGDSKRTAEAVAG
GS1 RREGATVIFVAVDGKLAGLIAIADPVKATTPAALAALRAAGLRIVMLTGDNRTTAEAVAR
GS2 RGEGATVMFVAVDGKLAGLIAVADPIKATTPAALKALHAAGLRIVMLTGDNRTTAEAVAR

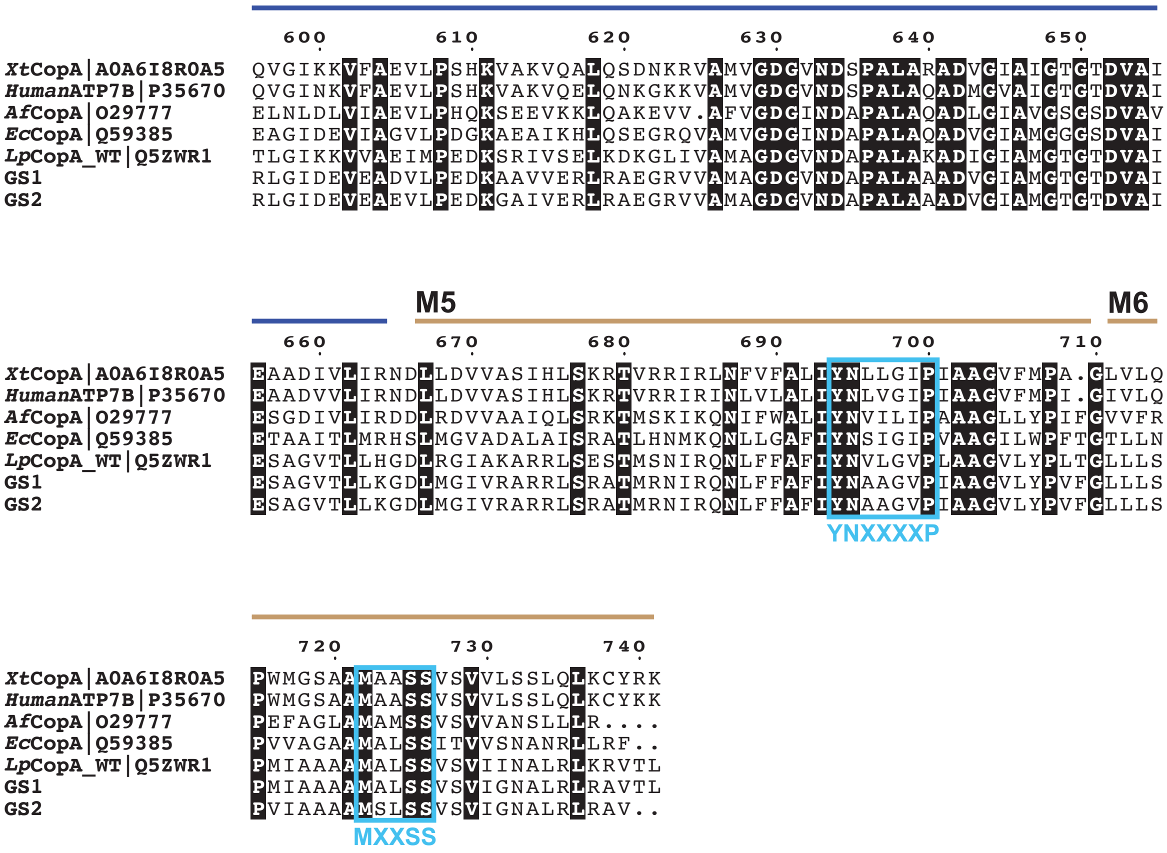


**Figure S3**: **Sequence alignment of various characterized** $P_{1B\text{-}1}$ **transporters with GS1 and GS2**. Domain boundaries and key motifs are labeled for clarity and comparison This larger set of sequences was aligned using the Clustal Omega tool hosted by EMBL-EBI, and the final figure was generated using ESPript3 (*62–64*).

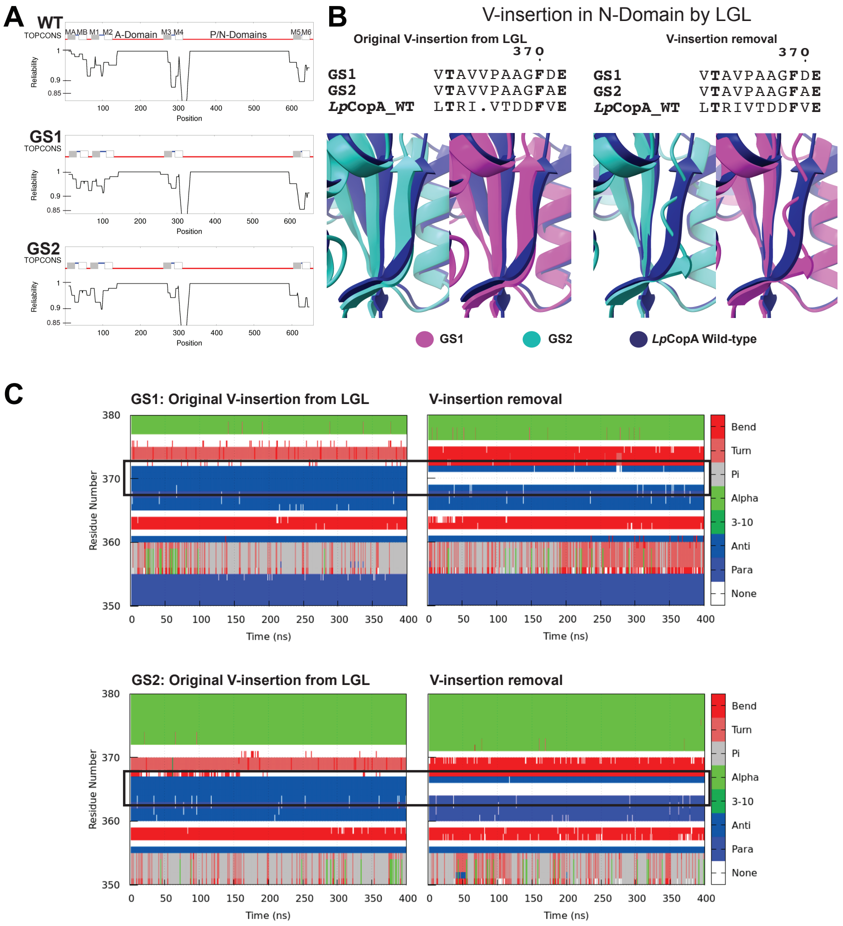


**Figure S4**: **Computational pre-screening and closer examination of the V-insertion in the N-domain of GS1 and GS2.** (**A**) TOPCONS, a tool for delineating intracellular and extracellular domains within a sequence (*28*), predicted a topology for both GS1 and GS2 that closely resembles that of WT *Lp*CopA, featuring eight transmembrane helices and two extensive intracellular regions. (**B**) AF2-predicted structures demonstrate that the V-insertion (valine) in both GS1 and GS2 is essential for the proper formation of the first N-domain $\beta$-sheet (P359 to A367 in GS1 and P358 to A366 in GS2) (*1*). (**C**) Secondary structure analysis indicates a failure to retrieve the corresponding $\beta$-sheet during simulations, as evidenced by the absence of secondary structure in the black box bounded region.

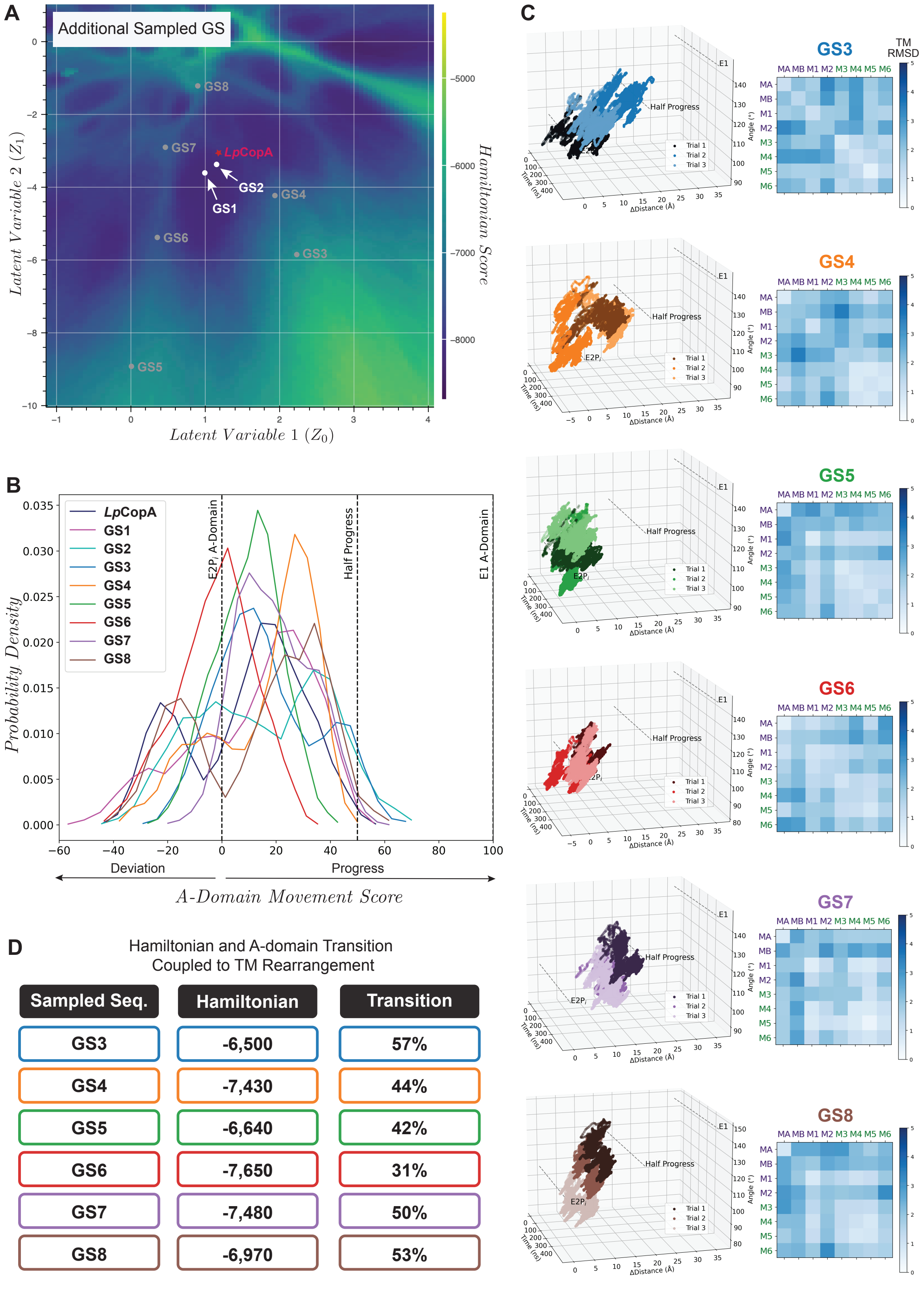


**Figure S5**: **MD analysis of additional generated sequences** (**A**) Decoding location of GS3-GS8. (**B**) Comparison of *A-domain movement score* reveals that most generated sequences fail to replicate the results of WT. (**C**) TM rearrangements were relatively weak, with two-block movements notably absent. (**D**) Hamiltonian values from LGL and the coupled A-domain transition progress corresponding to the reported TM rearrangement that is shown in part C.

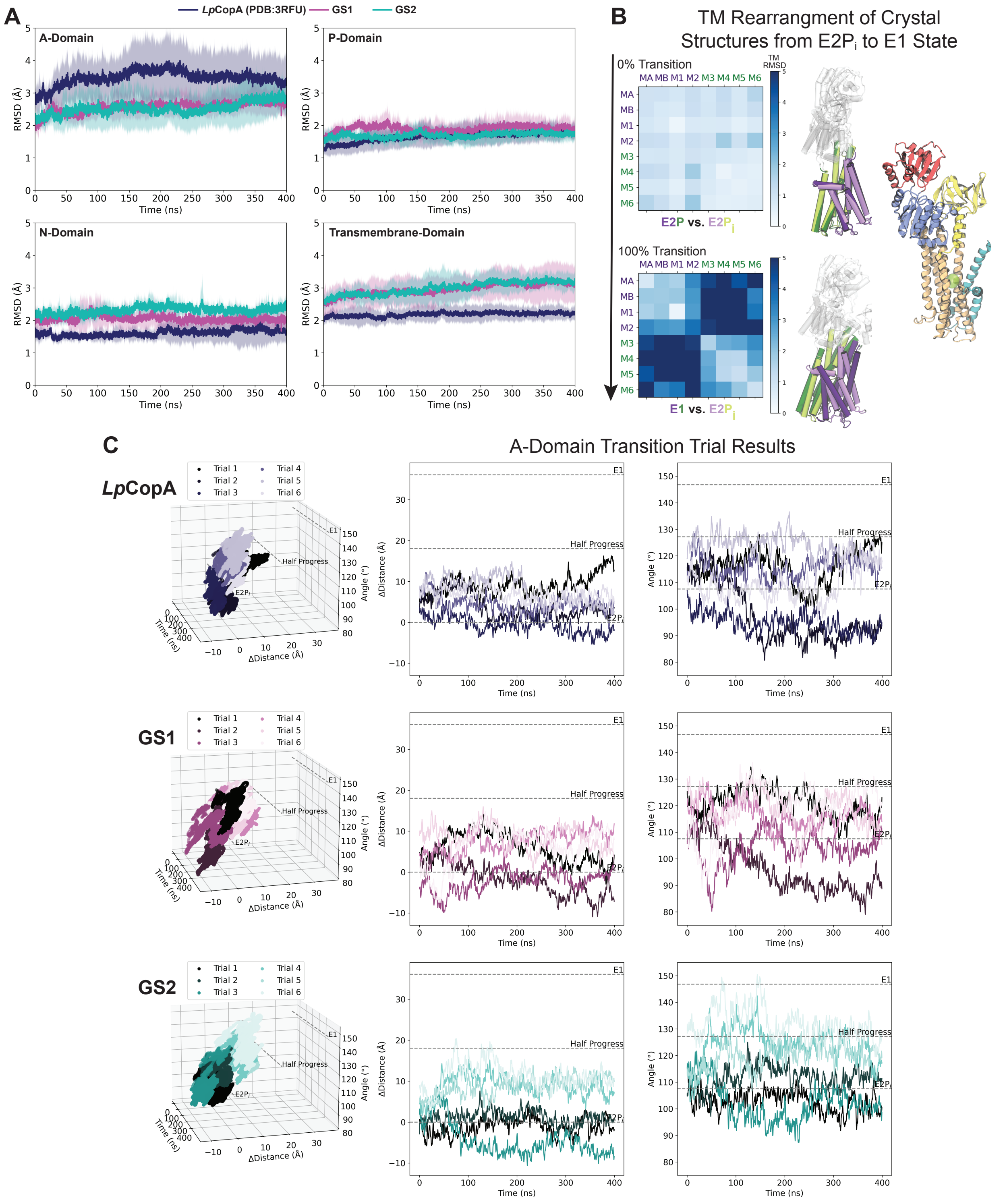


**Figure S6**: **Additional MD simulation details for *Lp*CopA, GS1 and GS2.** (**A**) Average RMSD comparisons of the different domains derived from MD simulations to monitor system stability. (**B**) Complete transmembrane rearrangement coupled to the A-domain transition, as determined from crystal structures of the E2$P_i$ (PDB: 3RFU; *Legionella pneumophila*, *Lp*CopA) and early-E1 (PDB: 7R0I; *Archaeoglobus fulgidus*, *Af*CopA) states. (**C**) Results from all six simulation trials that were utilized to determine the *A-domain movement score* from the E2$P_i$ to the E1 state in Fig. 3D.

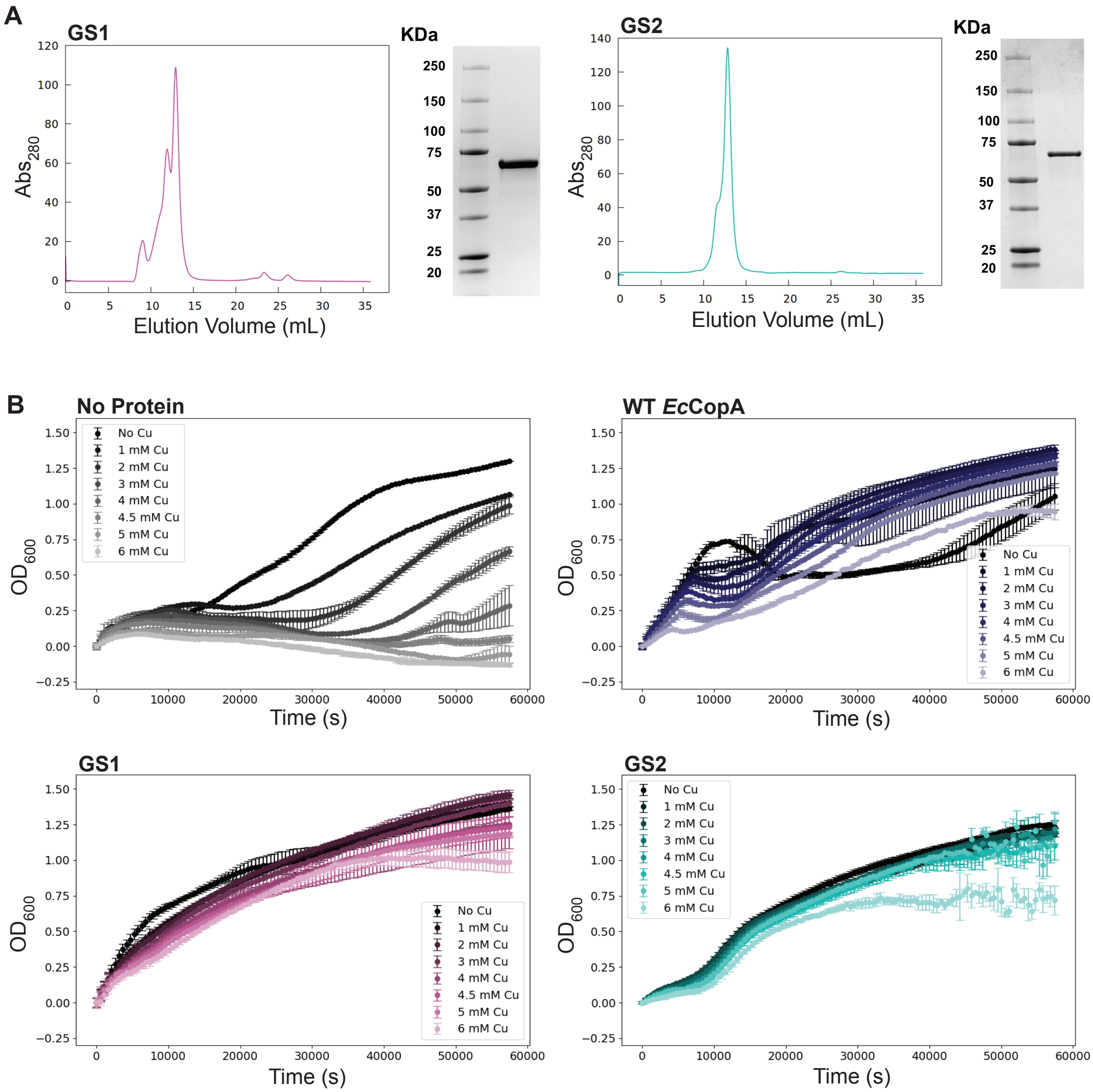


**Figure S7**: **Generated sequences purification and additional biochemical characterization.** (**A**) Size exclusion chromatography analysis of purified GS1 and GS2 with corresponding SDS-PAGE. (**B**) $OD_{600}$ results from a representative copper susceptibility growth assay

**Table S1. Maximum coupled *A-domain movement score* achieved across trials.**

| | Structure | Trial 1 | Trial 2 | Trial 3 | Trial 4 | Trial 5 | Trial 6 |
|---|---|---|---|---|---|---|---|
| *A-domain movement score* - Progress (%) | E2P (CS) | 0.00 | – | – | – | – | – |
| | E1 (CS) | 100.00 | – | – | – | – | – |
| | WT | 52.91 | 30.99 | 3.64 | 47.26 | 57.78 | 36.47 |
| | GS1 | 50.64 | 20.60 | 15.51 | 45.39 | 51.37 | 51.05 |
| | GS2 | 11.25 | 19.96 | 25.84 | 53.74 | 46.16 | 64.49 |
| | GS3 | 22.72 | 57.11 | 25.03 | – | – | – |
| | GS4 | 43.35 | 32.14 | 43.54 | – | – | – |
| | GS5 | 35.56 | 41.58 | 34.23 | – | – | – |
| | GS6 | 18.62 | 8.12 | 31.19 | – | – | – |
| | GS7 | 50.03 | 28.48 | 28.45 | – | – | – |
| | GS8 | 53.02 | 49.85 | 12.39 | – | – | – |

**Table S2. Maximum coupled inter-block helix DDM achieved across trials. Parenthetical values show the larger standard deviation between rows and columns in the inter-block DDM.**

| | Structure | Trial 1 | Trial 2 | Trial 3 | Trial 4 | Trial 5 | Trial 6 |
|---|---|---|---|---|---|---|---|
| Averaged Inter-block RMSD (Å) | E2P (CS) | 1.12 (0.38) | – | – | – | – | – |
| | E1 (CS) | 5.07 (0.51) | – | – | – | – | – |
| | WT | 1.93 (0.43) | 1.47 (0.45) | 1.57 (0.24) | 1.83 (0.38) | 1.82 (0.25) | 1.53 (0.49) |
| | GS1 | 1.35 (0.48) | 1.25 (0.24) | 1.13 (0.22) | 1.75 (0.41) | 2.19 (0.31) | 1.86 (0.62) |
| | GS2 | 1.02 (0.17) | 1.13 (0.33) | 1.33 (0.58) | 2.56 (0.75) | 2.75 (0.38) | 2.26 (0.66) |
| | GS3 | 1.99 (0.32) | 2.23 (0.46) | 2.13 (0.46) | – | – | – |
| | GS4 | 2.07 (0.72) | 2.37 (0.53) | 2.17 (0.43) | – | – | – |
| | GS5 | 1.61 (0.25) | 2.06 (0.18) | 1.75 (0.23) | – | – | – |
| | GS6 | 1.67 (0.41) | 1.42 (0.25) | 2.06 (0.37) | – | – | – |
| | GS7 | 1.78 (0.57) | 2.57 (1.02) | 2.03 (0.41) | – | – | – |
| | GS8 | 2.06 (0.29) | 1.72 (0.45) | 1.78 (0.39) | – | – | – |

**Supplementary Movie 1**. Representative simulation trial showcasing A-domain dynamics over 400 ns simulation by tracking changes in tilt angle and Δdistance for GS2.

**Supplementary Movie 2**. Representative simulation trial showcasing transmembrane helices rearrangement by tracking the Distance Difference Matrix over 400 ns simulation for GS2.
